# Paired guide RNA CRISPR-Cas9 screening for protein-coding genes and lncRNAs involved in transdifferentiation of human B-cells to macrophages

**DOI:** 10.1101/2021.04.26.441397

**Authors:** Sebastian Ullrich, Carme Arnan, Carlos Pulido-Quetglas, Ramil Nurtdinov, Alexandre Esteban, Joan Blanco-Fernandez, Estel Aparicio-Prat, Rory Johnson, Sílvia Pérez-Lluch, Roderic Guigó

## Abstract

CRISPR-Cas9 screening libraries have arisen as a powerful tool to identify both protein coding (pc) and non-coding genes playing a role along different processes. In particular, the usage of a nuclease active Cas9 coupled to a single gRNA has proven to efficiently impair the expression of pc-genes by generating deleterious frameshifts. Here, we first demonstrate that the usage of a second gRNA targeting the same gene synergistically enhances the capacity of the CRISPR-Cas9 system to knock out pc-genes. We next take advantage of our paired-guide (pgRNA) system to design a library to simultaneously target 874 pc-genes and 166 lncRNAs which are known to change expression during the transdifferentiation from pre-B cells to macrophages. We show that this system is able to identify known players in this process, and also predicts 26 potential novel ones, of which we select four for deeper characterization. Two of these, *FURIN* and *NFE2*, code for proteins related to cell differentiation and macrophage function; the other two, *LINC02432* and *MIR3945HG*, are lncRNAs associated with cancerous and infectious diseases, respectively. The CRISPR-Cas9 coupled to pgRNAs system is, therefore, a suitable tool to target simultaneously pc-genes and lncRNAs for genomic perturbation assays.

## INTRODUCTION

CRISPR-Cas9 library screening has become a powerful technique to identify genes, both protein coding genes (pc-genes) and long non-coding RNAs (lncRNAs), that play functional roles in cellular processes, such as cell differentiation or cancer progression. Actually, candidates identified in this type of screens have been proposed as potential therapeutic targets, reviewed in (1).

CRISPR screens targeting protein-coding genes (pc-genes) are mainly based on single guide RNAs (sgRNA) libraries that induce indel mutations in the target genes, leading to frameshifts and, consequently, loss of protein function, reviewed in (2–5). However, a recent study revealed residual protein activity for some targets after induction of frameshift mutations, leaving room for improvement (6). This sgRNA strategy is particularly ineffective when targeting long-non coding RNAs (lncRNAs), as point mutations or small indels will, in most cases, not affect the activity of the transcript. Thus, an alternative approach to impair lncRNA expression is to promote a deletion covering its promoter region and transcription start site (TSS) by means of a paired guide RNA (pgRNA) design. With this goal, we and others have developed different methods involving two guide RNAs cloned in the same vector, both targeting the same gene (7–9). Specifically with our system DECKO (Double Excision CRISPR Knockout) we were able to efficiently promote deletions of up to 3 Kb in cells expressing the Cas9 nuclease (7).

While the function of many lncRNAs remains unknown, few CRISPR “loss of function” screens have been performed targeting this gene class specifically. In one of these studies, the authors designed a paired guide RNA CRISPR-Cas9 library targeting 700 lncRNAs with potential oncogenic or tumor suppressor activity in human hepatoma cancer cells (10). From the initial candidates, only 7% of lncRNA were identified as positive hits, and only 1.3% were further validated. More recently, the same group developed a functional screening targeting splice sites of 10,996 multi-exonic lncRNAs in K562 cells. In this case, the authors identified 2% of lncRNAs essential for cell growth but only a small percentage of them could be further validated (11). In another work, an inactive Cas9 was fused to a transcription repressor, such as KRAB (CRISPR-inhibition, CRISPRi), which was used to knock down the expression of more than 16,000 lncRNAs. Almost 3% lncRNA involved in cellular growth were identified in a number of cell lines using this alternative approach (12). These studies highlight the high cell type specificity of many lncRNAs, which is especially convenient for tissue-specific targeted therapies. Of note, none of the aforementioned screenings were designed to target both coding and non-coding genes simultaneously.

Cell transdifferentiation is the process by which differentiated somatic cells are reprogrammed into other cell types without transitioning through a pluripotent state. Transdifferentiation is a powerful tool to effect the conversion from one cell type into another, being more efficient and less costly than conventional reprogramming and subsequent differentiation. This is of special interest for the development of novel therapies, such as the generation of cell types with specific features to promote regeneration after tissue injury or degeneration, reviewed in (13). Thus, the study of the genetic basis and the molecular changes occurring during transdifferentiation is essential to understand and control the conversion between cell types.

One powerful transdifferentiation model is the conversion of human B-cell precursor leukemia cells (BLaER1) to macrophages (14). BLaER1 cells are derived from the RCH-ACV lymphoblastic leukemia cell line (15). BLaER1 pre-B cells are able to transdifferentiate into macrophages upon induction in a process that lasts 7 days. These pre-B cells stably express the hematopoietic transcription factor ratCEBPa fused to an estrogen receptor (ER) hormone binding domain. When β-estradiol is added to the medium, it binds to CEBPaER and allows its translocation into the nucleus, where it induces the transcriptional program leading to macrophage morphology and function (14). During the transdifferentiation process, it is crucial to shut down the B-cell related expression program and activate the macrophage related one. However, the means by which CEBPa orchestrates the transdifferentiation process remains elusive.

With the goal of discovering pc-genes and lncRNAs that are essential for the transition from B-cell to macrophage, and taking advantage of available RNA-Seq data produced along the transdifferentiation process (16), we have used the DECKO system with a combined library of paired guide RNAs (pgRNAS), targeting simultaneously 166 lncRNAs and 874 pc-genes upregulated along the transdifferentiation process. Towards that end, we have extended the CRISPETa bioinformatics pipeline (17) to design optimal pairs of sgRNAs for deletion of genomic regions including both pc-genes and lncRNAs. We have observed that targeting pc-genes with two gRNAs synergistically enhances the CRISPR knockout efficiency. The results from our screen suggest that the transdifferentiation from B-cell into macrophage is very robust, and very few genes are able to perturb the progression of the process. Still, out of the targeted genes, we identified 26 candidate genes potentially delaying the transdifferentiation, seven of which were individually validated. Among them, two pc-genes, *FURIN* and *NFE2*, and two lncRNAs, *LINC02432* and *MIR3945HG*, were further interrogated at genomic and transcriptomic level, confirming the efficiency of the pgRNA DECKO system in knocking out protein and lncRNA expression. *FURIN* and *NFE2* have been previously involved in myeloid branch differentiation (18–20) and *MIR3945HG* has been found overexpressed in tuberculosis infected human macrophages (21). This confirms that, with our system, we have indeed been able to specifically uncover both pc-genes and lncRNAs involved in blood differentiation.

## MATERIALS AND METHODS

### Target gene selection from transcriptomics data

The selection of target genes was based on RNA-seq data sampled at 12 time points (0h, 3h, 6h, 9h, 12h, 18h, 24h, 36h, 48h, 72h, 120h, 168h) during transdifferentiation of human BLaER1 cells to macrophages (16). The RNAseq data was quantified with GRAPE-nf (https://github.com/guigolab/grape-nf). Read mapping was performed with STAR (22) and gene expression quantification with RSEM (23) using the GENCODE annotation v22 (24). Two biological replicates were analyzed separately.

The 19,814 pc-genes and 14,855 lncRNAs (union of the following biotypes: processed transcript, 3 prime overlapping ncRNA, sense intronic, antisense, macro lncRNA, lincRNA, non-coding and sense overlapping from GENCODE v22) were filtered for a minimum average expression of at least 1 FPKM for pc-genes (0.1 FPKM for lncRNAs) and at least 4x fold change for protein pc-genes (2x fold for lncRNAs) between highest and lowest expression value along the temporal profile. In addition, lncRNAs were required to have a minimum expression of 1 FPKM in at least one time point and to be non overlapping with other genes in a 5 Kb window on the same strand and 50 bp on the opposite strand relative to their TSS. This resulted in 4,804 pc-genes remaining for replicate 1 and 4,552 for replicate 2, and 642 lncRNAs for replicate 1 and 536 for replicate 2. Those genes were clustered separately for each replicate into 36 expression profiles for pc-genes and 16 for lncRNAs with k-means clustering in R. We focused on two types of expression profiles: “peaking profile” (genes that increase their expression level at the beginning of the transdifferentiation process and later on decrease) and “upregulated profile” (genes that are upregulated throughout the process). Pooling those profiles within each replicate and then intersecting between the replicates, resulted in a final list of 939 protein-coding and 174 lncRNA candidate genes.

### Paired guide RNA library design

For lncRNAs, CRISPETa (17) was used to target genes’ TSS. For pc-genes, we developed a new version of CRISPETa to target ORFs (code available at https://github.com/Carlospq/CRISPETa_PC). In this case, we first obtained the principal isoform from the APPRIS database (25). The exonic sequence of this isoform was extracted from the human genome sequence version h19, using the GENCODE annotation v22, and searched for all possible protospacers (20mers followed by a PAM sequence of NGG). sgRNA were scored using the RuleSet2 algorithm (26) and paired. Pairs were ranked according to: 1) location in the ORF sequence, 2) the pair score calculated as the sum of the two individual sgRNA scores, and 3) the deletion region of the pair (prioritizing those predicted to create an out-of-frame deletion). The first coding exon was preferentially targeted. In case not all designs could be placed at the first coding exon, the window was extended to the second and third exons. For lncRNAs, the region targeted around the TSS was increased stepwise from 500 to 5,000 bp in consecutive runs of CRISPETa until the required number of pgRNAs was designed. Selected pgRNAs for lncRNAs were filtered so as to not overlap pc-genes. In all cases, sgRNAs were filtered to remove possible off-targets using CRISPETa’s pre-computed database with default value [-t 0,0,0,x,x] for the first run and relaxing this cutoff for consecutive runs, as described in (17). CRISPETa output parameters were adjusted to provide the sequence of the 165 nt oligonucleotide (Insert-1) needed for library cloning using DECKO method (7), which includes the targeting regions of the pgRNAs separated by a cloning site (Supplementary Table S2).

Up to ten pgRNAs were designed per target gene with a minimum distance of 50 bps between any pair of gRNAs. In total, we designed pgRNAs for 166 lncRNAs and 874 pc-genes. In addition, we designed 50 pgRNAs for each ratCEBPa, SPI1 and ITGAM positive controls. For negative controls, we designed pgRNAs for 100 intergenic regions, 10 pgRNAs each. As a non-targeting negative control for library sorting assays we used a pgRNA against Firefly luciferase, called “pDECKO-non targeting”.

### Library cloning

A ssDNA library of 12,000 oligos of 165 nt (insert-1) (Supplementary Table S2) was purchased from Twist Biosciences. The library was amplified to obtain dsDNA using emulsion PCR as described in (27), and cloned into pDECKO_mCherry vector ((17), Addgene 78534) following the 2 cloning steps described in (7). ENDURA electrocompetent cells (Bionova Cientifica) were used to ensure high efficiency transformation and avoid recombination errors. Several transformations were performed in parallel. For the first cloning step (intermediate plasmid), approximately 500,000 bacterial colonies were collected and processed together in a single maxiprep. To eliminate the remaining empty plasmid, we took advantage of the fact that insert-1 (in the intermediate plasmid) contains unique restriction sites (EcoRI and BamHI), which are not present in the original backbone. Digesting the intermediate plasmid resulted in a linear product that could be distinguished from the circular empty backbone and purified in an agarose gel. For the 2nd step of cloning, 50 ng of BsmbI-digested intermediate plasmid was mixed with 1 μl annealed Insert-2 (gRNA1 constant region coupled to an H1 promoter, previously assembled from four oligonucleotides and diluted 1:20) and 1 μl of T4 DNA ligase (Thermo Scientific) and incubated for 4h at 22ºC (as described in (7)). Several transformations with ENDURA electrocompetent cells were done in parallel. For the 2nd cloning step (final plasmid) more than 100,000 bacterial colonies were collected and processed together in a maxiprep. A scheme of the final plasmid can be found in Supplementary Figure S4A.

### Cell culture and library infection

Human BLaER1 cells (14) were kindly provided by Thomas Graf (CRG, Barcelona) and grown in RPMI medium (Invitrogen) supplemented with 10% heat-inactivated foetal bovine serum (FBS), 2 mM L-glutamine, and 100 U/ml penicillin G sodium (14). BLaER1 cells were first infected with a plasmid containing Cas9 fused to BFP ((17), Addgene 78545), selected for more than 5 days with blasticidin (15 μg/ml) and sorted using a BD FACS Aria instrument at the Flow Cytometry Unit of the Center for Genomic Regulation. These cells, stably expressing Cas9, were then infected with the pDECKO library. For lentivirus production, we performed 80 co-transfections of HeK293T virus packaging cells (at approximatelly 60-70% confluence on 10 cm dishes) with 3 μg of the pDECKO_mCherry plasmid library and 2.25 μg of the packaging plasmid pVsVg (Addgene 8484) and 750 ng of psPAX2 (Addgene 12260) using Lipofectamine 2000 (according to manufacturer’s protocol). Transfection media was changed on the following day to RPMI. In total, 400 ml of viral supernatant were collected 48h post transfection, filtered through a cellulose acetate filter, and used for overnight infection of 90×10E6 BLaER1-Cas9 cells at a density of 250,000 cells/ml with presence of polybrene (10 μg/ml). The percentage of infection was computed as the number of mCherry positive cells compared to the total number of cells with a Fortesa cell cytometer analyser. Infection rate ranged between 2-4%, ensuring a low multiplicity of infection (less than 1 viral integration per cell) (28). After 48h of infection, the cells were double selected with blasticidin (20 μg/ml) and puromycin (2 μg/ml) for 18-19 days. 15 million of the BLaER1-Cas9 library infected cells were induced for transdifferentiation into macrophages by using β-estradiol, IL-3 and M-CSF, as described previously (14). After incubation for 3 days (T3) /6 days (T6) they were collected for FACS sorting.

### Individual target validation

For paired guide RNA pDECKO-mCherry plasmid cloning we used the method described in (17) (sgRNA sequences are listed in Supplementary Table S1 and the cloning oligos are detailed in Supplementary Table S3). For single guide RNA pDECKO-mCherry plasmid cloning we used the method described in (29) (see Supplementary Table S4 for details of the oligos used). Plasmids constructed for this study can be found in Supplementary Table S5 (plasmids available at Addgene.org are indicated).

For lentivirus production, we co-transfected HeK293T virus packaging cells with 3 μg of each pDECKO_mCherry plasmid and packaging plasmids as described previously. Viral supernatant was collected 48h post transfection and filtered through a cellulose acetate syringe filter. Polybrene (10 μg/ml) was added. We pelleted 5×10E5 BLaER1-Cas9 cells in two microcentrifuge tubes and resuspended each of them with 1 ml of viral supernatant. We performed spin-infection for 3h at 1,000 g. After infection, the viral supernatant was removed and infected cells were resuspended with RPMI media supplemented with 10% heat-inactivated foetal bovine serum (FBS), 2 mM L-glutamine, and 100 U/ml Penicillin Streptomycin. After 48h of infection, we performed double selection with blasticidin (20 μg/ml) and puromycin (2 μg/ml) antibiotics. The selection was maintained for a minimum of 2 weeks.

### Flow cytometry

#### For cell sorting

30×10E6 cells were counted and resuspended in 300 μl PBS + 3% FBS in the presence of FcR blocking reagent. Cells were incubated for 10 minutes and 15 μl of the human anti-CD19 antibody conjugated with BV510 (Becton Dickinson, 562947) and 15 μl of human anti-cd11b (Mac-1) antibody conjugated with PE-Cy7 (Labclinics, 25-0118-41) were added. Cells were incubated for 30 minutes in the dark, washed with PBS and resuspended in 2 ml of PBS + 3% FBS. Topro-3 was added as a viability marker. Cells were sorted in a BD FACS Aria instrument at the Flow Cytometry Unit of the Center for Genomic Regulation.

#### For flow cytometry analysis

1×10E6 cells were counted and resuspended in 100 μl PBS + 3% FBS in the presence of FcR blocking reagent. Cells were incubated for 10 minutes and 5 μl of each of the corresponding antibodies were added. For the CD19 knockout experiment, we used the antibody anti-CD19 conjugated with PE-Cy7 (Becton Dickinson, 557791). Cells were incubated for 30 minutes in the dark, washed with PBS and resuspended in 500 ul of PBS + 3% FBS. Topro-3 was added as a viability marker. Cells were measured in a BD Fortessa analyser. For the Stain Index calculation we used the formula: (mean positive - mean background) / (2 * SD background), as previously described (30).

### Sample processing for deep sequencing

Genomic DNA was extracted from the FACS sorted cells with the GeneJET Genomic DNA purification kit (Thermo Scientific) and 2 PCR steps were performed. A scheme of oligo binding sites is shown in Supplementary Figure S4.

A first PCR step was done by Phusion polymerase (Thermo Fisher) using 500 ng of genomic DNA and staggered oligo mix (Supplementary Table S6) with the presence of 6% DMSO, annealing temperature of 60ºC and a total of 20 cycles of amplification. We used staggered oligos to avoid the same bases being read for the constant region during Illumina sequencing and to minimize technical issues during base calling. Up to 6 PCR reactions were combined, the amplicons were gel-purified, and 2 ng were used as a template for a second PCR.

The second PCR step was also done by Phusion polymerase but without the presence of DMSO. We used Illumina barcoded oligos (Supplementary Table S7), an annealing temperature of 60°C and a total of 8 cycles of amplification. Samples were purified with Agencourt Ampure beads (Beckman Coulter), quantified with a Qubit fluorometer (Thermo Scientific) and checked for quality in a Bioanalyzer (Agilent). We then pooled the libraries and sequenced them on the Illumina HiSeq 2500 at the Genomics Unit of the Center for Genomic Regulation (150 bp paired-end sequencing) to have about 20 million reads per sorted subfraction.

### Mapping and quantification of sequencing reads

For read mapping, based on the initial pgRNA library with two guides per target (Supplementary Table S2), an artificial genome was generated by concatenating the 41 bp of the two pgRNAs (gRNA1 21 bp, gRNA2 20bp) and converted into FASTA format. STAR mapper (version 2.4.2a) (22) was used to index the genome, adjusting the standard settings by the following parameter for small genomes:

--genomeSAindexNbases 6

In the resulting genome after removing duplicated constructs, each pgRNA pair is represented by each one of the 11,550 chromosomes with a length of 41 bp.

Dynamic trimming of Illumina reads was done in perl by pattern matching the insertion site of the pgRNAs in the plasmid sequence (“ACCG” for pgRNA1 in the window of 15-55 bp of read2, “AAAC” for pgRNA2 in the window of 100-150 bp of read1). The extracted 20 bp fastq sequences for the pgRNA2 were reverse complemented and concatenated to the 21 bp fastq sequences for the pgRNA1. Fusion reads with fewer than 20 bp sequence length were filtered out.

Mapping was performed with STAR version 2.4.2a with the following parameters:

STAR --runMode alignReads --runThreadN 8 --genomeDir /users/resources/genome -- readFilesCommand zcat --readFilesIn pgRNA1_pgRNA2.fastq.gz --alignIntronMax 1 --outSAMtype BAM SortedByCoordinate --outSAMunmapped Within --limitBAMsortRAM 3000000000 --outFilterMultimapNmax 1 --outFilterMismatchNmax 11 --outFilterMatchNmin 30 --outFilterMatchNminOverLread 0.1 --outFilterMismatchNoverLmax 0.9 --outFilterScoreMinOverLread 0.1

Given the distance between the sequencing primer and gRNA2, the pipeline was conceived to be adjustable to a variable number of mismatches. Running the pipeline without allowing for any mismatches, we could only make use of about 25 to 30% of the reads. Hence, we increased the number of allowed mismatches in progressive steps that resulted in a steep increase of mapped reads until a saturation point was reached between 10-15 mismatches, depending on the sample (Supplementary Figure S6C). For further analysis, we allowed for a maximum of 13 mismatches to stay below 1% of multi-mapped reads for all samples of both replicates. Spearman correlation values of 0.95-1.00 between samples, mapped with zero mismatches compared with up to 13 mismatches, justified the usage of the quantification data with substantially more reads and therefore higher statistical power (Supplementary Figure S6D). For quantification, the count for each guide pair within the mapped libraries was aggregated from the BAM files with SAMtools (31).

Due to the low memory footprint of the artificial genome, this quantification strategy can be applied even on laptops with moderate specifications (minimum requirements: single core CPU, 4GB RAM, 10GB disk space).

The mapped reads were clustered to check for reproducibility between replicates (data not shown).

### LNA GapmeRs assay

LNA antisense oligonucleotide GapmeRs (Exiqon) complementary to human lncRNA *LINC02432* (ENSG00000248810.1) (GCATGAAAGAGTTGGT) and lncRNA *MIR3945HG* (ENSG00000251230.1) (CTGAGAGGTGGCAAGC) were designed. A LNA oligonucleotide containing a scrambled sequence (AACACGTCTATACGC) was used as a negative control. We seeded 40,000 BLaER1 cells in a 24-well plate and the cells were grown in 1 ml complete RPMI media containing LNA GapmeRs at a final concentration of 2 μM. After 3 days of incubation, we induced transdifferentiation as described previously (14). Total RNA was isolated from cells after 3 days of induction.

### RNA extraction, retro-transcription and quantitative PCR

RNA extractions from 1×10E6 cells were performed with Quick RNA Miniprep Kit (Zymo Research). 140 ng-500 ng RNA were retro-transcribed with Reverse Aid reverse transcriptase (Thermo Scientific). Quantitative PCR (qPCR) was performed with NZY Speedy qPCR Green Master mix (NZY tech) and in a LightCycler 480 Real-Time PCR System (Roche). Primer sequences are detailed in the Supplementary Table S8. Quantifications were normalized to an endogenous control (Glyceraldehyde 3-phosphate dehydrogenase, GAPDH). The relative quantification value for each target gene compared with the calibrator is expressed as 2^(Ct-Cc).

### Western blot

1×10E6 cells were resuspended with 100 μL of Lysis buffer (1% SDS, 10 mM EDTA, 50 mM Tris pH 8, protease inhibitors). The cell lysate was sonicated in a Branson sonicator for 10 seconds (50% amplitude and power 7). The samples were run in a 10% SDS-PAGE gel and transferred to a nitrocellulose membrane. The membrane was blocked with blocking buffer (TBS, 0.1% Tween 20, 5% non fat milk) O/N at 4°C, and incubated for 1h 30’ at room temperature with primary antibodies: anti-FURIN rabbit polyclonal antibody (Proteintech, 18413-1-AP) 1:1,000 in blocking buffer or anti-NFE2 rabbit polyclonal antibody (Proteintech, 11089-1-AP) 1:1,000 in blocking buffer. After 5 washes with TBS-0.1% Tween 20, the membranes were incubated for 1h with the secondary antibody goat anti-rabbit-HRP (Sigma, G9545) 1:10,000 in blocking buffer. After 5 washes with TBS-0.1% Tween 20, the membranes were incubated either with Amersham ECL western blotting detection reagent (GE Healthcare, RPN2209), or Super Signal West Femto Maximum Sensitivity Substrate (Thermo Fisher, 34096), and imaged in an Amersham Imager 600. As a protein loading control, the membranes were re-blotted with primary antibody rabbit anti-GAPDH-HRP polyclonal antibody (Proteintech, 10494-1-AP) 1:4,000 in blocking buffer, and incubated for 1h at room temperature. Washes and secondary antibody incubation were performed as previously described. The presence of two bands in NFE2 western blot likely corresponds to different post-translational modifications of NFE2 (18).

### TA cloning

In order to sequence the edited region in BLaER1-Cas9 cells, we amplified the deletion junctions by PCR using oligos outside the cut region (Supplementary Table S9). The resulting PCR products were cloned using a TA cloning kit (Life Technologies), according to manufacturer’s instructions, and sequenced by Sanger sequencing.

## RESULTS

### Cellular model and target selection

BLaER1 is a leukemia B cell line able to transdifferentiate into macrophages through the stable expression of the ratCEBPa transcription factor fused to an estrogen receptor hormone binding domain (14) (Figure 1A). During transdifferentiation, the changes in the cell identity can be monitored by flow cytometry through the tracking of specific cell surface markers. For example, the expression of CD19, found on B-cells, decreases during the process until disappearing, not being detected in the transdifferentiated macrophages, whereas Mac-1, a macrophage surface marker, starts appearing in the transdifferentiating B-cells at 36 hours after induction and its detection is maximized at the end of the process (Figure 1B).

**Figure 1:**
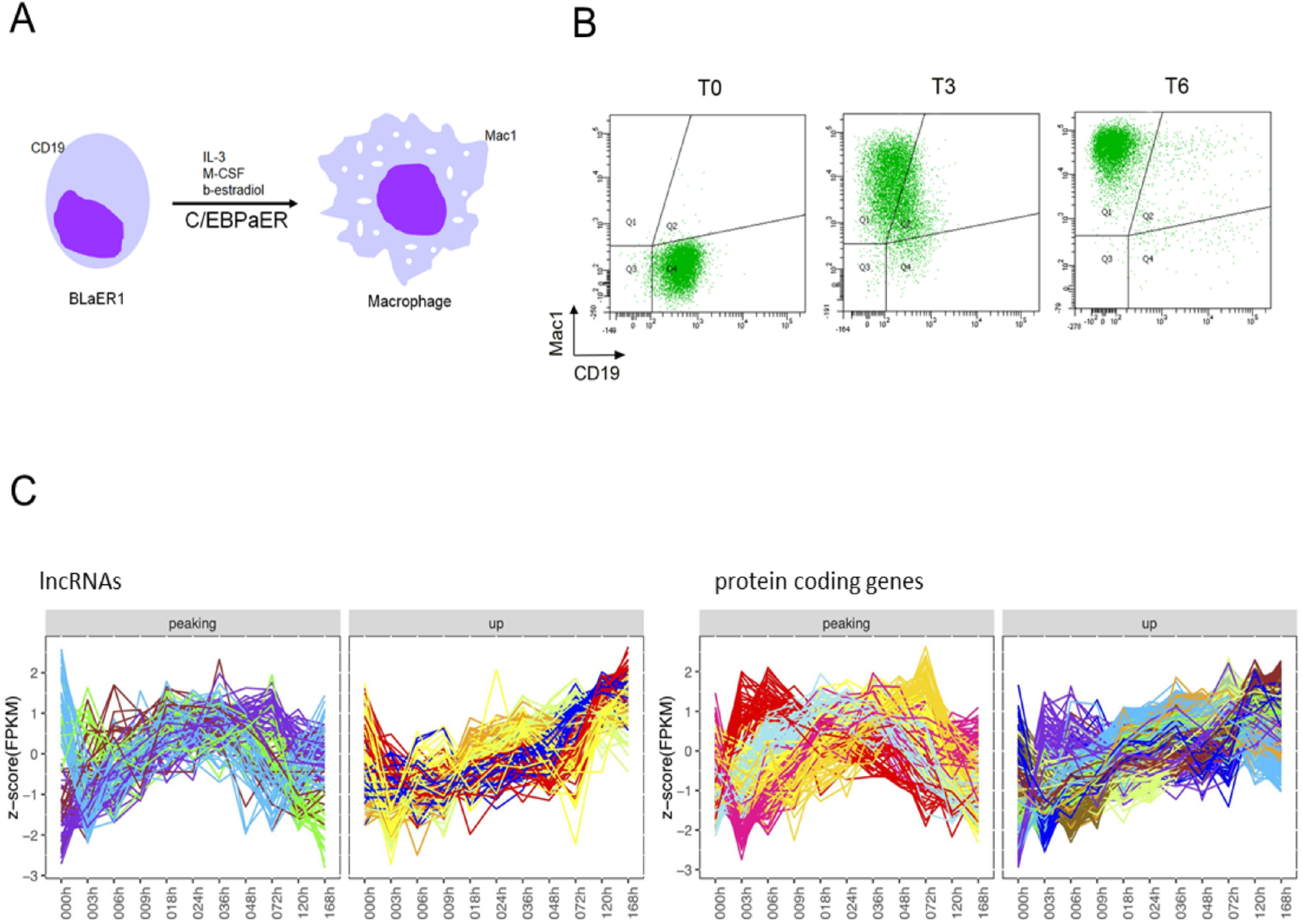
Cellular model and targets selection. (**A**) Transdifferentiation of BLaER1 pre-B cells into macrophages is accompanied by a dynamic transcriptomic remodelling of the cells. BLaER1 lymphocytes transdifferentiate into functional macrophages in the presence of Interleukin 3 (IL-3) and Macrophage colony-stimulating factor (M-CSF) upon β-estradiol induced release of CEBPaER to the nucleus. (**B**) Flow cytometry analysis of cell surface markers at T0, T3 and T6 after induced transdifferentiation in BLaER1-Cas9 cell line. During the process, BLaER1 cells progressively lose the CD19 (B-cell marker staining -X-axis-) and gain the Mac-1 (macrophage marker staining -Y-axis-). (**C**) Merged k-means clustered expression profiles (color code) of peaking and upregulated genes during transdifferentiation: 16 initial clusters of lncRNA (n=174) and 36 initial clusters of protein coding genes (n=939). FPKM values were log10 transformed before the normalization to z-score. Each line shows the expression pattern of a gene along transdifferentiation. The color corresponds to the k-means cluster to which the gene belongs (see also Supplementary Figures S1 and S2).

To identify coding and non-coding genes that may drive the BLaER1 transdifferentiation process, we analyzed available RNA-seq data at 12 time points along the seven days the process lasts, in two biological replicates (16). We identified 488 lncRNAs and 3,627 pc-genes with minimum expression values higher than 1 FPKM in at least one time point as well as expression changes higher than 2-fold for lncRNAs and 4-fold for pc-genes (see Methods). We clustered the 4,115 genes with k-means into 16 lncRNA and 36 protein coding clusters (Supplementary Figures S1 and S2). After visual inspection, genes from all clusters showing upregulated and peaking profiles were selected as candidates to be involved in the transdifferentiation process, comprising in total 174 lncRNAs and 939 pc-genes (Figure 1C). For both lncRNAs and pc-genes, upregulated genes showed higher expression than peaking genes, which peaked at about 36 hours (Supplementary Figure S3).

### A CRISPR knockout library targeting simultaneously non-coding and protein coding genes

We first asked whether a pgRNA format could yield improved rates of knockout for pc-genes compared to sgRNAs. Towards that end, we designed a set of gRNAs aganist the lymphocyte B surface marker CD19 and infected Cas9 expressing BlaER1 cells with either individual gRNAs or pgRNAs (Supplementary Tables S1 and Figure 2A, upper panel). In order to quantify the efficiency of the knockout, we collected the infected cells and stained them with a fluorescently conjugated anti-CD19 antibody. Single gRNAs caused a 30% to 70% decrease of CD19 immunofluorescence with the only exception of construct CD19-4, where the single gRNAs caused a decrease ranging from 83% to 97% when compared to the negative control (Figure 2A, lower panel). Although the knockout efficiency of single gRNAs is very variable, the decrease in CD19 signal is enhanced when the cells were infected with any combination of pgRNA (approximately 96% signal reduction), indicating that the effect of using more than one gRNA per target gene is more than additive.

**Figure 2:**
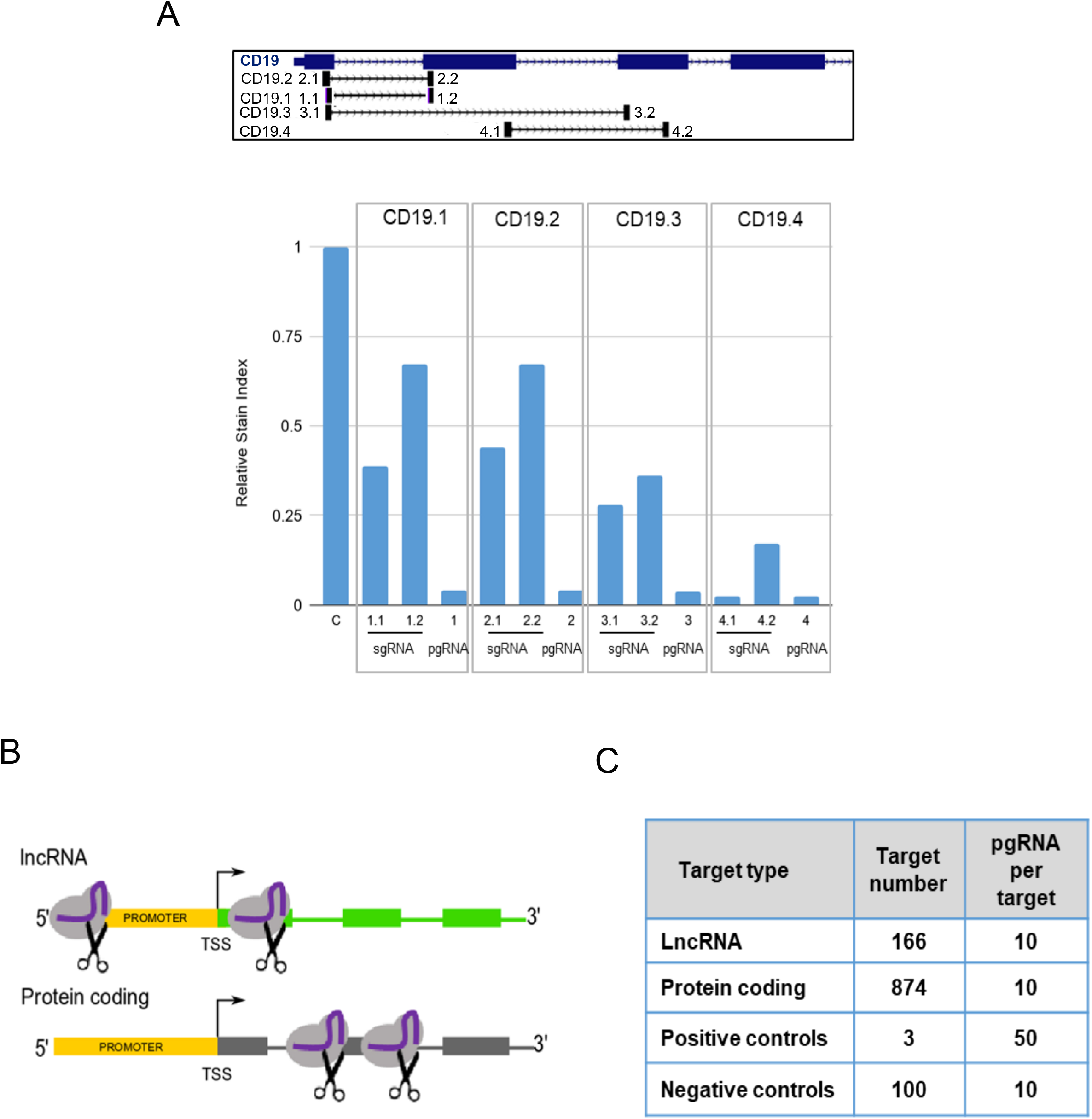
pgRNA CRISPR library for lncRNA and pc-genes. (**A**) (Upper panel) Diagram of the CD19 gene indicating the target sequence of CD19 pgRNAs (sgRNA1 and sgRNA2, from left to right). (Lower panel) Flow cytometry analysis of fluorescence intensity of the CD19 protein in BLaER1-Cas9 cells infected with sgRNAs and pgRNAs. The relative Stain Index of the different infected cells compared to the maximum expression level of CD19 in control cells (BLaER1-Cas9 cells infected with pDECKO-GFP (7)) is represented. CD19 expression is reduced between 30% and 95% upon infection of sgRNAs. The infection of pgRNAs induces a consistent reduction of CD19 signal up to 95% with all pgRNAs tested. (**B**) Schematic diagram showing the position of pgRNAs targeting lncRNAs (targeting the promoter and the transcription start site) and pc-genes (targeting coding exons). (**C**) CRISPR library composition (number of targets of each biotype and pgRNA pairs designed per target).

With the goal of uncovering which peaking and upregulated genes are necessary for the progression of the transdifferentiation, and given the strong synergistic effect observed when targeting pc-genes with two gRNAs, we designed a combined pgRNA CRISPR library targeting simultaneously the coding exons of the pc-genes and the promoter and the TSS region of the lncRNAs (Figure 2B) identified above (see methods, Supplementary Figure S4, and Supplementary Table S2). Using CRISPETa (17), we designed a CRISPR library targeting the 174 lncRNAs. In parallel, we developed a new version of CRISPETa (see Methods) to specifically target protein coding genes, and used it to design pgRNAs targeting the 939 pc-genes selected above at a depth of 10 unique pgRNAs each. According to our on- and off-target filters, we managed to design pgRNAs targeting the TSS of 166 lncRNAs and the ORFs of 874 pc-genes (see Methods, Figure 2C). As controls, pgRNAs targeting pc-genes necessary for transdifferentiation, namely ratCEBPa - a transcription factor used to induce the transdifferentiation (14) -, SPI1 - a downstream transcription factor activated by CEBPa needed for both B cell and macrophage differentiation (32,33)-, and ITGAM - a subunit of the Mac1 complex used to track macrophage differentiation-(positive controls), and 100 intergenic regions (negative controls) were added to the library. The CRISPETa output including the pgRNA generated oligonucleotides can be found in Supplementary Table S2.

### CRISPR-Cas9 screen of genes required for transdifferentiation

To identify the genes involved in the transdifferentiation from B cells to macrophages, BLaER1-Cas9 cells were infected with the combined library at low multiplicity of infection (Figure 3A). In parallel, a plasmid containing non-targeting pgRNAs was transduced as negative control. Cells were collected at 3 days (T3) and 6 days (T6) after induction, and their transdifferentiation status was tracked by flow cytometry with B-cell and macrophage-specific markers (Figure 3B and Supplementary Figure S5). We expected to find pgRNAs targeting genes required for transdifferentiation in the “delayed” cell population, that progresses at a slower rate compared to the control cells (quadrant Q4, corresponding to undifferentiated cells, in Figure 3B compare left -control- vs right -library- panels and Supplementary Figure S5).

**Figure 3:**
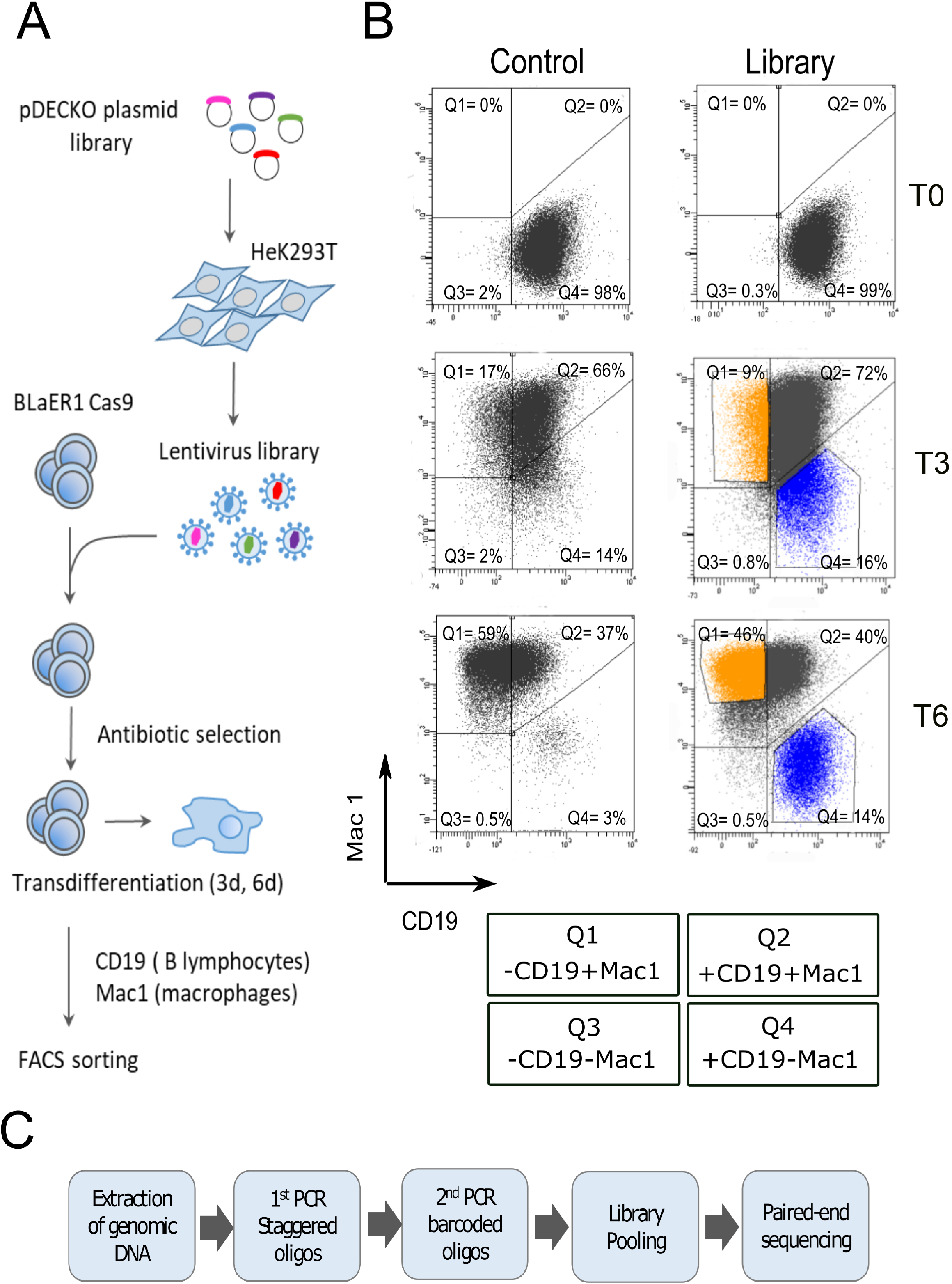
CRISPR-Cas9 screening in BLaER cells. (**A**) Workflow of the CRISPR screening experiment. The pDECKO plasmid library was transfected into HeK293T cells to obtain a library of lentivirus. BLaER1-Cas9 cells were infected at low multiplicity of infection and double selected with antibiotics (Blasticidin and Puromycin) for 20 days. The infected cells were induced for transdifferentiation into macrophages for 3 days (T3) and 6 days (T6). Cells were labelled with antibodies against cell surface markers: CD19 (for B-lymphocytes) and Mac-1 (for macrophages). Transdifferentiation status was assessed by flow cytometry. Transdifferentiated and delayed populations were isolated by Fluorescence-Activated Cell Sorting (FACS). (**B**) Flow cytometry analysis of BLaER1-Cas9 cells infected with the pDECKO_non-targeting control (left panels) and with the pDECKO_CRISPR-library (right panels) at T0, T3 and T6 of transdifferentiation. CD19 antibody, conjugated with BV510 fluorophore, was used to identify B-cells and Mac-1 antibody, conjugated with PE-Cy7 fluorophore, was used to identify macrophages. Quadrants are as follows: Q1 (macrophage-like cells with presence of Mac-1 and absence of CD19 surface markers); Q2 (transition cells with the presence of Mac-1 and CD19); Q3 (background and not stained cells, negative for Mac-1 and CD19); Q4 (lymphocyte B-like cells with the presence of CD19 and absence of Mac-1 surface markers). The percentage of cells in each of the 4 quadrants is shown. The fraction of sorted cells showing a delay of transdifferentiation (“delayed” fraction) is marked in blue (gate P4), and sorted cells that differentiate at a normal pace (“differentiated” fraction) is marked in orange (gate P5). See also Supplementary Figure S5. (**C**) Workflow of the processing of the sorted cell populations for deep sequencing. Genomic DNA of sorted cells was extracted and PCR amplified in 2 steps. For the 1st PCR, specific staggered primers were used to amplify the integrated fragment which contains the pgRNAs. For the 2nd PCR, Illumina barcoded primers were used to allow for sample pooling. Samples were sequenced by 150 bp paired-end Illumina sequencing. See also Supplementary Figure S4.

Whereas the library infected cells only showed a mild delay in comparison to the negative control at T3 of transdifferentiation (16% vs 14% in Q4, respectively) (Figure 3B and Supplementary Figure S5), the difference was much stronger at T6 (14% vs 3% in Q4, respectively). To identify the pgRNAs responsible for the delay of transdifferentiation, the delayed (blue gates) and the differentiating (orange gates) populations were recovered by fluorescence activated cell sorting (FACS) at T3 and T6 after transdifferentiation induction, and the genome integrated pgRNAs were sequenced (see Methods).

### Identification of lncRNAs and protein coding genes involved in delay transdifferentiation

On average, 25 million reads were sequenced for each isolated population. We implemented a bioinformatics protocol to analyze and quantify these reads (Supplementary Figure S6, see Methods). The distribution of pgRNAs of the original library after cloning showed a similar profile to the distribution of pgRNAs identified upon transdifferentiation induction (T0), demonstrating that all pgRNAs in the initial library were represented in the screening. However, during the course of the transdifferentiation (T3 and T6), a small fraction of guide pairs became enriched while many others were depleted (Supplementary Figure S7).

To identify the pgRNAs enriched in each subpopulation of cells, we defined the differentiation delaying effect (DDE) as the ratio of counts of a given pgRNA from the delayed subpopulation (del) divided by the counts from the transdifferentiated population (dif). Thus, larger DDE values represent genes required for the correct transdifferentiation. DDE was computed independently for the two replicates at T3 and T6 after transdifferentiation.

We first assessed whether the DDE score could distinguish between positive and negative controls. Indeed, ratCEBPa pgRNAs showed reproducible large DDE values that correlates between replicates (for all tested sets of pgRNAs, Figure 4A left panel). The values for all the intergenic pgRNAs were much lower (Figure 4A right panel), showing no reproducibility between replicates.

**Figure 4:**
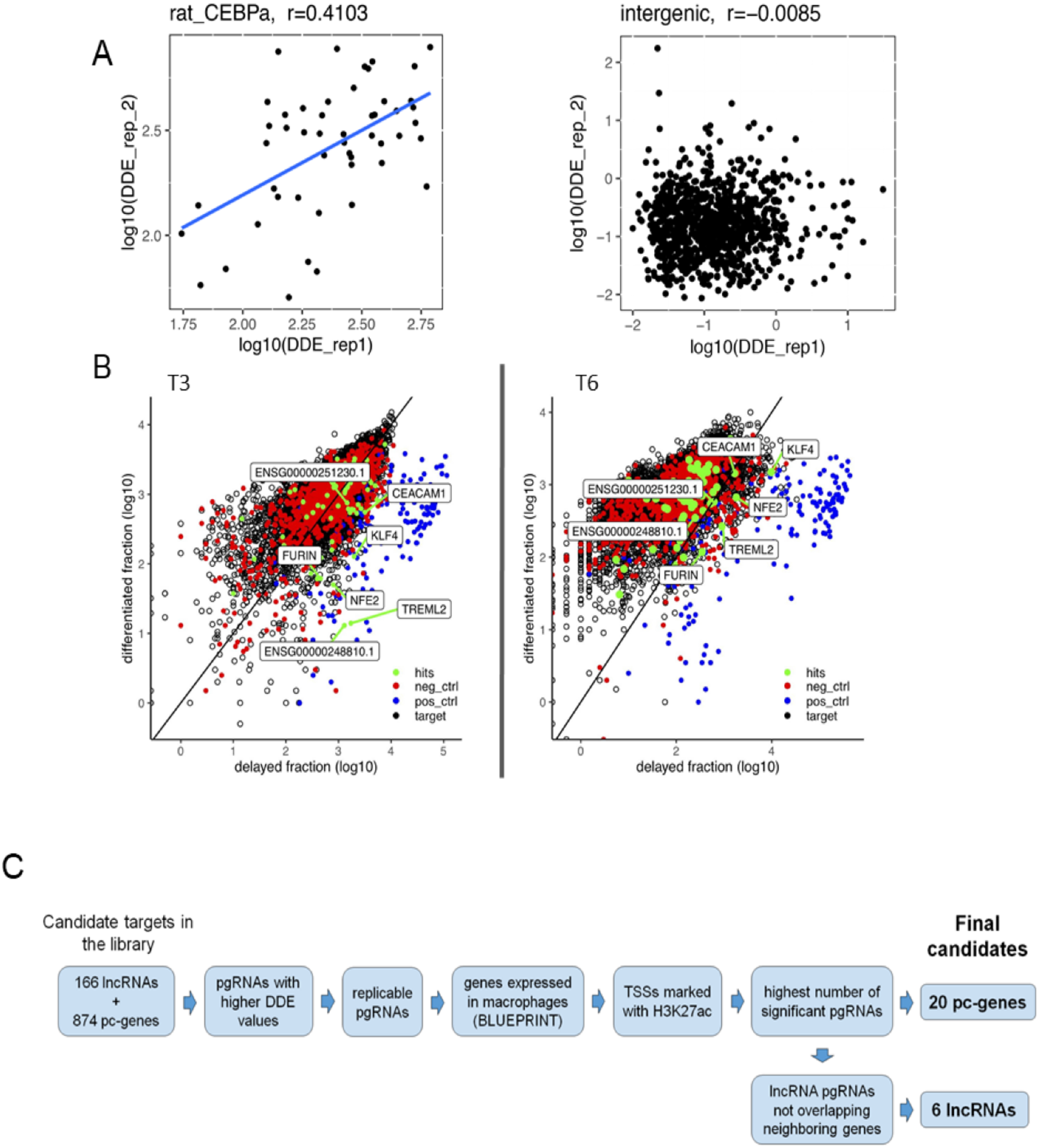
Identification of lncRNAs and protein coding genes involved in transdifferentiation. (**A**) Correlation between replicates of the differentiation delaying effect (DDE, ratio of reads from delayed versus transdifferentiated fraction) observed per pgRNA of ratCEBPa (left panel) and intergenic negative controls (right panel) after 6 days (T6) of transdifferentiation. Spearman correlation values are stated above. The DDE of CEBPa pgRNAs are very large and show a positive correlation between replicates, whereas intergenic pgRNAs do not show reproducible DDE between replicates.(**B**) Scatterplot of log10 transformed counts in delayed versus differentiated fractions at T3 and T6 after induction of transdifferentiation. pgRNAs targeting positive controls are depicted in blue, non-targeting pgRNAs in red, screened candidates in black, and selected hits for validation in green. Final candidate genes are also highlighted. ENSEMBL genes ENSG000002448810.1 and ENSG00000251230.1 correspond to *LINC02432* and *MIR3945HG* lncRNAs respectively. (**C**) Decision tree followed to identify candidate genes, from the CRISPR-Cas9 screening, involved in the transdifferentiation process. From the original list of 1,040 pc-genes and lncRNAs, we ended up with a set of seven candidates to undergo further validation.

We next plotted the count distribution of the pgRNAs detected in the delayed fraction against the differentiated fraction (Figure 4B). Confirming the efficiency of the methodology, pgRNAs targeting positive controls (blue) showed a higher enrichment in the delayed fraction compared to the differentiated one, whereas negative control pgRNAs (red) moved from the diagonal at T3 to the differentiated fraction at T6. Although, at T3, the bulk of pgRNAs targeting candidate genes were centered around the diagonal, a number of pgRNAs showed enrichment in the delayed population, which was attenuated with an overall shift of guides towards the differentiated fraction at T6 of transdifferentiation.

In order to identify potential target genes affecting the transdifferentiation process, we selected all pgRNAs with DDE values in the highest decile (at T3 DDE > 1.89, at T6 DDE > 0.44, mean of both replicates) (Supplementary Figure S8A). Besides that, for T3 and T6 separately, we required potential targets to have at least two identical pgRNA pairs in the upper decile for both biological replicates. Following this criteria, 18 lncRNAs and 86 pc-genes were selected at T3 and 50 lncRNAs and 135 pc-genes at T6 time point. The union of candidates from both time points resulted in a total of 64 lncRNAs and 191 pc-genes (Supplementary Table S10). Comparing the distribution of the DDE values of all the pgRNAs corresponding to the selected target genes against positive and negative controls revealed significant differences between them, especially at T3 after induction, when both lncRNA and protein coding targets show significantly higher DDE values than the negative intergenic controls (Supplementary Figure S8B).

To further narrow down the candidate list for individual validations, we applied additional criteria (Figure 4C). First, we checked the consistency of the expression of the candidates along the hematopoietic tree (Blueprint RNA-seq quantifications from the Blueprint Dataportal http://dcc.blueprint-epigenome.eu/) and discarded candidates with either no expression in B-cells/macrophages or unexpected relative expression, e.g. significantly lower expression in macrophages than in B-cells. Second, we selected candidates that showed H3K27ac marking at the TSS along seven ENCODE cell lines, a mark that has been related to both active promoters and enhancers (34,35). Third, we selected the candidate genes with the highest number of pgRNAs significantly enriched in the delayed population in comparison to the differentiated cells. Finally, for lncRNAs, we further verified that the pgRNAs targeting the promoter region did not overlap any other neighboring gene. Considering all these criteria, we ended up with 6 lncRNAs and 20 pc-genes as candidates to be involved in transdifferentiation. From them, the top two lncRNAs -*LINC02432 and MIR3945HG* - and five protein coding genes - *FURIN, NFE2, KLF4, TREML2* and *CEACAM1*-, following the aforementioned criteria, were selected as the targets with the highest potential to impact transdifferentiation efficiency (RNA expression profiles of these candidates along transdifferentiation can be found in Supplementary Table S11).

We next wanted to assess if the candidate genes identified in the CRISPR screening did, indeed, play a role during the transdifferentiation process. First, we individually validated the delay of positive control pgRNAs, such as CEBPa and SPI1, compared to the intergenic negative ones. The delay effect was measured by tracking the expression of CD19 and Mac-1 at T3 and T6 after induction (delayed cells are represented in the Q4 quadrant, Figure 5A). Indeed, we observed a strong delay for cells expressing pgRNA against CEBPa and SPI1 compared to intergenic regions both at T3 and T6 (Figure 5) indicating that the efficiency of the pgRNA knockout is very high when targeting protein coding genes, which is consistent with the high efficiency also observed when knocking out CD19 (Figure 2A).

**Figure 5:**
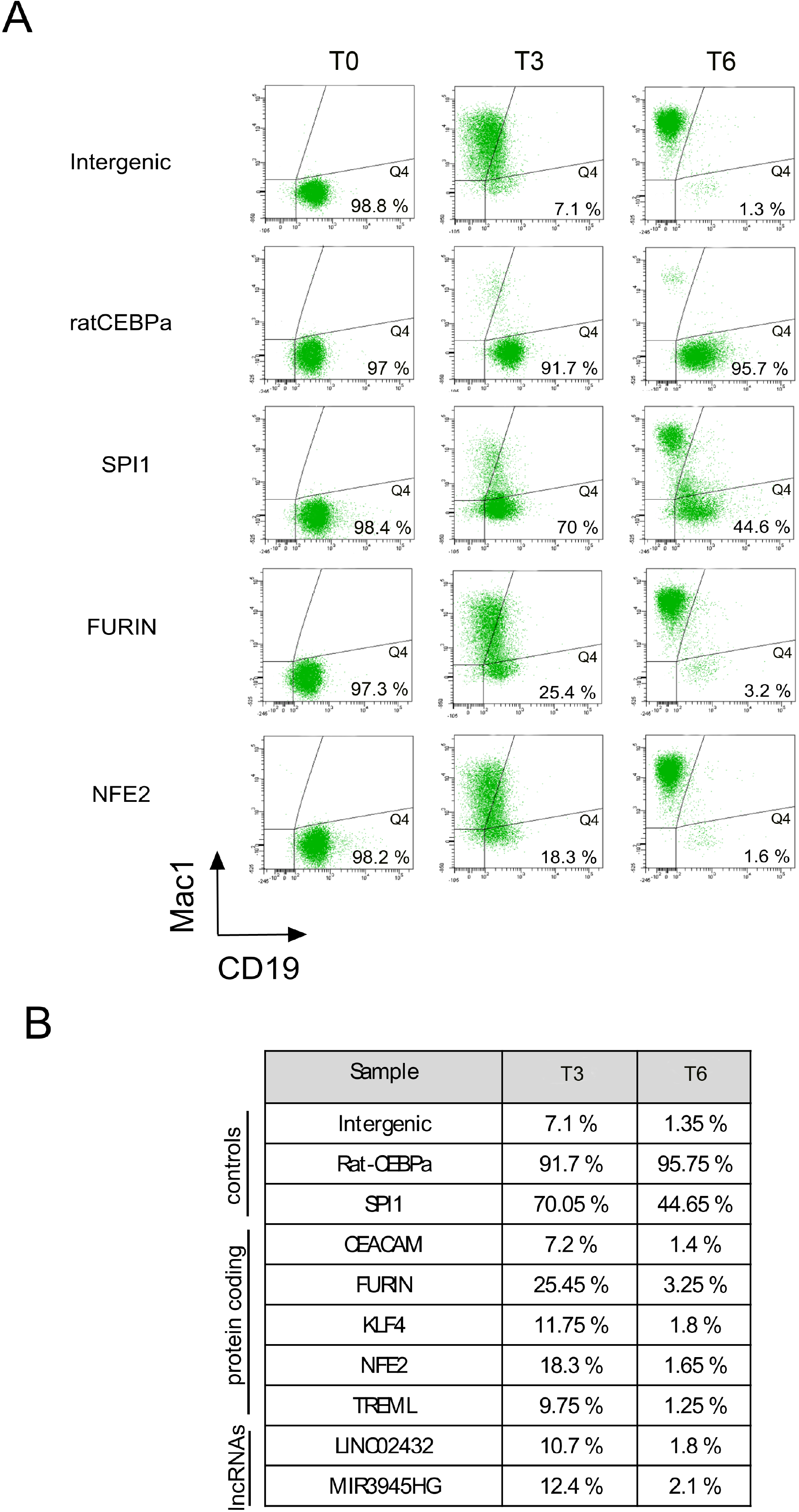
Individual target validation by flow cytometry. Flow cytometry analysis of control and candidate pgRNAs at T0, T3 and T6 after induction of transdifferentiation. (**A**) Flow cytometry plots of intergenic negative control, two positive controls targeting ratCEBPa and SPI1, and two protein coding targets *FURIN* and *NFE2*. CD19 B-cell marker is represented on the X-axis and Mac-1 macrophage marker is represented on the Y-axis. Cells that do not undergo transdifferentiation remain in the Q4 quadrant (positive for CD19 -X-axis- and negative for Mac-1 -Y-axis) (the percentages of cells in this quadrant is shown). (**B**) Percentage of cells with delayed transdifferentiation (Q4 quadrant) observed in controls and individually validated candidates (average of 2 biological replicates) at T3 and T6 after induction of transdifferentiation.

Next, we selected the pgRNAs showing the highest enrichment in the library for each of the five protein coding and the two lncRNA candidate genes (Supplementary Table S1), and infected BLaER1-Cas9 cells. Cells infected with pgRNAs against the two lncRNAs (*LINC02432* and *MIR3945HG*) showed some initial delay in transdifferentiation (10-12% at T3), whereas they seem to fully recover at T6, showing delays comparable to the negative intergenic controls (Figure 5B and Supplementary Figure S9 A-B). For pc-genes, the knockout of *FURIN* and *NFE2* had the strongest delaying effect on transdifferentiation (25% and 18% at T3 respectively, Figure 5). For the remaining genes tested, undifferentiated cells ranged between 10 and 12%, except for *CEACAM1*, which showed a delay comparable to the intergenic negative control (Figure 5B and Supplementary Figure S9 C-G).

### Individual validation of lncRNAs involved in the transdifferentiation process

To further validate the role of the two candidate lncRNAs in transdifferentiation, we next assessed whether the CRISPR-Cas9 was able to efficiently induce a deletion at the promoter region of these genes. Thus, we FACS isolated the cell populations corresponding to the delayed fractions from the individual validations above and amplified and sequenced the region surrounding their TSS. Indeed, we could validate the double cut, as well as diverse rearrangements with multiple indels, in the clones from the *LINC02432* targeted cells tested (Supplementary Figure S10 A-B). In the case of the *MIR3945HG*, however, we were not able to confirm the deletion (data not shown).

To distinguish if the role of these lncRNAs on transdifferentiation was mediated by the RNAs themselves or by a putative enhancer effect of the DNA regions transcribing the lncRNAs, we designed LNA GapmeRs against the two lncRNAs targeting the same isoforms as were depleted by CRISPR (Supplementary Figure S10 C-D). Although the expression of both lncRNAs was significantly impaired upon GapmeR treatment, we did not observe any transdifferentiation delay. This suggests that the impact of the deletion of these two lncRNAs on the process is likely due the disruption of a possible enhancer activity of the genomic sequence. Consistent with this hypothesis, both loci are enriched in H3K27ac, mark associated to active enhancers (34), and this enrichment is enhanced upon induction of the process (Supplementary Figure S11) (16).

On the other hand, we also analyzed recently released data on active/silent compartments, A/B compartments, during BLaER1 cells transdifferentiation (36). We found that *MIR3945HG* stays in the A compartment along the process. *LINC02432,* instead, is found in the B, inactive, compartment in pre-B cells, but at 72 hours it turns into A compartment, turning inactive again later in transdifferentiation, reflecting the peaking expression profile of the gene (Supplementary Table S11). We believe that this behavior further supports the implication of this lncRNA in the transdifferentiation process.

### Individual validation of protein coding genes involved in the transdifferentiation process

Regarding the pc-gene candidates, we validated *FURIN* and *NFE2* at genomic, transpriptomic and protein level. At genomic level, we could identify different editing events (indels) at the *FURIN* locus (Supplementary Figure S12A-B). Note that some of the clones do not show the long deletion expected if both pgRNAs induced the Cas9 cut. Still, targeting only one of the two regions can generate a frameshift, resulting in a non-functional protein. At transcriptomic level, *FURIN* expression, measured by qRT-PCR, decreased to around 50% in the full population of infected cells at T3 compared to the intergenic negative control (Figure 6A, FUT3 vs. CT3, respectively). This decrease reaches 70% when only the delayed population is measured (Figure 6A, FUT3s). Although we do not expect the deletion of an internal part of the gene to affect transcript abundance, we hypothesize that the lack of functional protein may cause the degradation of the transcript by nonsense mediated decay. We also observed that the decreased gene expression has an impact at protein level, as the FURIN protein is not detectable by western blot in CRISPR-Cas9 edited cells at T3, compared to the intergenic control, where a weak band is detected (Figure 6A).

**Figure 6:**
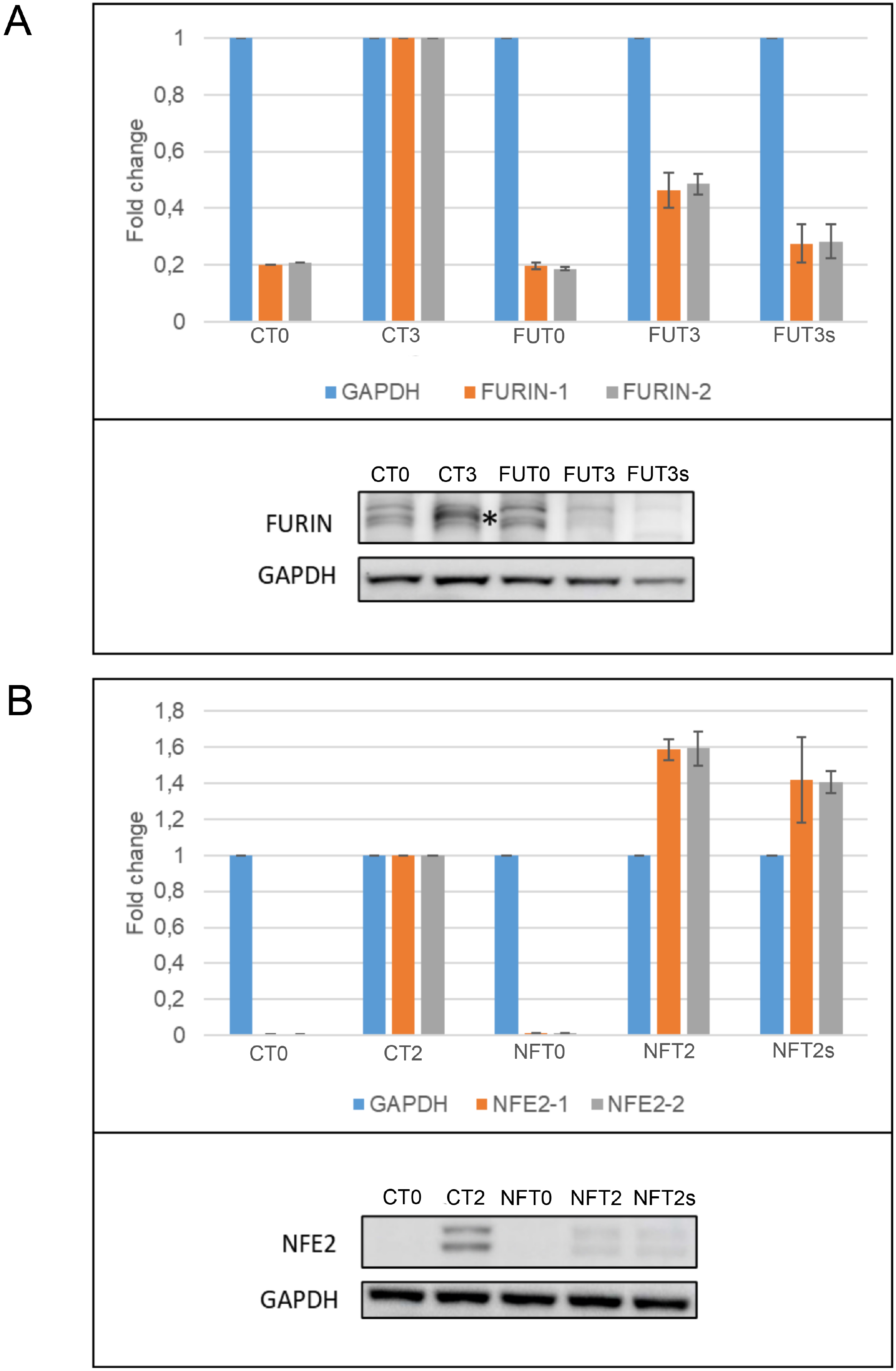
FURIN and NFE2 expression after CRISPR edition. (**A**) FURIN RNA and protein expression. Cells were collected at T0 (before induction) and T3 (3 days after transdifferentiation induction) . (CT0) and (CT3) negative control pDECKO-Intergenic at T0 and T3 respectively, (FUT0) and (FUT3) pDECKO-FURIN at T0 and T3, (FUT3s) pDECKO-FURIN at T3 and sorted from gate P4 (delayed population). Upper panel, qRT-PCR to check the expression of FURIN using two different sets of primers. Results are normalized to GAPDH and the fold change is calculated relative to the expression of cells infected with pDECKO-intergenic pgRNA at T3. The expression of FURIN decreases in cells infected with FURIN pgRNAs, especially in the delayed subpopulation (FUT3s). Bottom panel, western blot to assess the levels of the FURIN protein in BLaER1-Cas9 infected cells. Anti-FURIN antibody recognizes a band (marked with an asterisk), the signal of which increases at T3, in line with RNA-Seq data (Supplementary Table S11). The FURIN band is not detectable in the pDECKO-FURIN infected cells (FUT3 and FUT3s). (**B**) NFE2 RNA and protein expression. (CT0) and (CT2) negative control pDECKO-Intergenic at T0 (before induction) and T2 (2 days after transdifferentiation induction) respectively, (NFT0) and (NFT2) FU) pDECKO-NFE2 at T0 and T2, (NFT2s) pDECKO-NFE2 at T2 and sorted from gate P4 (delayed population). Upper panel, qRT-PCR to check the expression of NFE2 using 2 different sets of primers. Results are normalized to GAPDH and the fold change is calculated relative to the expression of cells infected with pDECKO-intergenic T2. NFE2 expression in NFE2 pgRNA targeted cells is higher than in intergenic control cells (NFT2 and NFT2s compared to Ct2). Bottom panel, western blot to check the protein levels of NFE2 in BLaER1-Cas9 infected cells. Anti-NFE2 antibody detects two bands, the signal of which increases at T2 (CT2 compared to CT0). These two bands are strongly reduced in NFE2 targeted populations (NFT2 and NFT2s compared to CT2).

For *NFE2*, we also identified small indels at genomic level (Supplementary Figure 12 C-D). In this case, RNA expression analysis showed that, for CRISPR-Cas9 edited cells, *NFE2* expression increases compared to negative control cells after 2 days of transdifferentiation (T2) (Figure 6B). Protein levels, in contrast, appear to decrease at this time point (Figure 6B). We hypothesize that this contrasting pattern between *NFE2* transcript and protein expression could potentially be explained by the production of non-functional protein promoting the continuous overexpression of the gene, in an attempt to overcome the lack of functional NFE2 protein.

## DISCUSSION

Along this manuscript, we have described the use of the CRISPR-Cas9 technology to identify genes, both lncRNAs and protein coding, involved in the transdifferentiation from pre-B cells into macrophages. With this goal we have designed the first library, to our knowledge, targeting simultaneously the TSS of lncRNAs and the coding region of pc-genes. We think that a combined library is a suitable approach to identify large numbers of target candidates independently of their biotype. Besides, our library design can be customized to target not only genes but also genomic regions putatively involved in dynamic processes, for instance enhancers or chromatin insulators.

We had already demonstrated that the DECKO system is able to efficiently induce deletions of up to 3 Kb around the TSS of lncRNAs, and that these deletions impaired gene expression (7). Here, we further wanted to assess whether using paired gRNAs would significantly increase the efficiency of the knockout of protein coding genes. Indeed, we found that targeting the CD19 B lymphocyte marker with paired gRNAs was more efficient than targeting it with a single gRNA. Actually, the decrease in protein expression after pgRNA infection was more than additive, meaning that the usage of two gRNAs is synergistically enhancing the efficiency of the CRISPR system. Consistent with this strong effect, we have also observed a strong transdifferentiation delay in cells infected with pgRNAs targeting *CEBPa* and *SPI1*. We hypothesize that the different efficiency observed between CEBPa and SPI1 knockdown may be due to differences in gene copy number. Although BLaER1 cells showed high levels of CEBPa compared to other clones (14), we speculate that *ratCEBPa* is only present at one copy per cell, as more than 90% of cells show a delay of transdifferentiation, indicating that almost all cells have been knocked out. In contrast, pgRNAs against *SPI1* are expected to target the two endogenous copies of the gene; thus, the fact that 45% of cells targeted with pgRNAs against *SPI1* show delayed transdifferentiation suggests that only in around 50% of cells the two *SPI1* copies have been efficiently knocked out. Overall, and given the high knockout rates observed in all cases, we think that the usage of two gRNAs can represent a convenient approach to target protein coding regions.

As a result of our screen we identified six lncRNAs and twenty protein coding genes as potential candidates to play a role in the transdifferentiation of B-cells to macrophages. One of the critical points in the library design (especially for lncRNAs) is the correct annotation of TSS (37), which is constantly improved and updated in the new GENCODE releases (38). The incomplete annotation of the non-coding genes may also influence the correct targeting of these genes, and may partially explain the relative low validation rate for lncRNAs in this type of screenings.

Our results demonstrate the high knockout efficiency of the DECKO system. Indeed, the 50 pgRNAs designed against CEBPa were positively enriched in the fraction of delayed cells, also highlighting the very good performance of CRISPETa predicting the pgRNAs with higher cutting scores (17). We think that the relatively low number of positive candidates may obey to the strong capability of BLaER1 cells to transdifferentiate. Accordingly, whereas most leukemia and lymphoma cell lines tested were not able to transdifferentiate in an efficient manner, almost 100% of BLaER1 cells were able to efficiently undergo the process, likely due to the constant and high expression levels of transgenic CEBPa (14). We think that the depletion of many factors involved in the transdifferentiation process cannot overcome the severe transcriptomic change induced by CEBPa in these leukemia B-like cells. Indeed, even when the knockout of some factors, such as FURIN and NFE2, is able to promote an initial delay of transdifferentiation in a high percentage of cells after induction (at T3), the targeted cells are able to eventually overcome the lack of the proteins and differentiate into macrophages after 6 days. The fact that different candidate genes were identified in the screening only at T3 or only at T6 after induction (no target appears enriched at both time points) also reflects the capability of BLaER1 cells to overcome the lack of these genes.

We observed a difference between the strength of the delaying effect of protein coding and lncRNA candidate genes. Whereas protein coding genes show between 10% -TREML2- and 25% -FURIN- delayed cells at T3 after induction, the two identified lncRNAs present 10-12% delayed cells. The stronger delaying phenotype observed for protein coding genes could be partially due to the high efficiency of one or the two pgRNAs to induce small genome rearrangements in the coding region, compared to lncRNAs, whose depletion implies the simultaneous action of both gRNAs and the induction of a deletion. Actually, it has been shown that the induction of CRISPR deletions is much less efficient (39), likely reducing the number of knockout cells.

Among the validated lncRNAs, *LINC02432* was previously identified as an upregulated lincRNAs in neuroblastoma cell lines (40) and *MIR3945HG*, which is overexpressed in macrophages upon infection with *Mycobacterium tuberculosis*, has been proposed as a candidate marker for the diagnosis of tuberculosis (21). The pc-genes had a comparable stronger effect. Among them, *FURIN* appears to play the stronger role. This protein is a ubiquitously expressed serine protease enzyme that processes substrates like cytokines, hormones, receptors and growth factors like TGFB1, which controls proliferation and differentiation in many cell types (41); and has been involved in tumour progression, representing an interesting therapeutic target. FURIN has been also related to monocyte/macrophage migration and proliferation, being also an inhibitor of apoptosis (19,20). Actually, the expression pattern of *FURIN* suggests that it is involved in the last steps of macrophage lineage determination, consistent with its role in macrophage motility. Another protein with notable effect is the transcription factor NFE2. This factor was found to be essential for regulating erythroid and megakaryocytic maturation and differentiation, but also impacts the renewal of hematopoietic stem cells (18,42,43). Altered NFE2 activity predisposed to leukemic transformation (44) and the NFE2 protein is overexpressed in the majority of patients with myeloproliferative neoplasms (45). Also the other protein candidates are related to blood and/or differentiation functions, although they showed milder effects when specifically validated. For instance, the transcription factor KLF4 is one of the Yamanaka factors and allows the induction of pluripotent stem cells from differentiated cells (46,47) and interacts with CEBPB, that is an important transcription factor that regulates the expression of genes involved in immune and inflammatory responses (48). TREML2 is a cell surface receptor that enhances T-cell activation, but it is expressed throughout the hematopoietic lineage (49,50) and it is coexpressed with TREML1 and TREM1, that stimulate monocyte-mediated inflammatory responses (51). Finally, the cell-cell adhesion molecule CEACAM1 has regulatory functions in T-cells, it is potentially important for the adhesion of macrophages at sites of infection (52), and it has also been associated to B cell aggregation in central nervous system autoimmunity (53). The fact that all these factors have been previously related to differentiation processes and/or blood associated functions indicate that, indeed, they are necessary for macrophage function or to induce differentiation of the myeloid branch, suggesting that they may be also necessary to promote BLaER1 pre-B cell transdifferentiation into macrophages.

All in all, we have designed a CRISPR-Cas9 library to simultaneously target lncRNAs and protein coding genes, and assess their role in B-cell to macrophage transdifferentiation. This screening has led to the identification of a few candidates that could potentially play a role in this process. The low number of candidates and the rapid recovery of the cellular perturbations induced by the CRISPR-Cas9 knockouts indicates, however, that the transdifferentiation of the BLaER1 cells into macrophages is a very stable and robust process. Nevertheless, we have demonstrated that the DECKO libraries are very efficient in promoting both frameshifts and deletions. We believe, therefore, that this is a powerful method for the study of the regulation of dynamic processes, as it is suitable for the deletion of small genomic regions, not only lncRNA TSSs, but also putative enhancers and other regulatory regions, as well as for the efficient knockout of protein coding genes.

## ACKNOWLEDGEMENTS

The authors would like to thank the CRG/UPF Flow Cytometry Unit for assistance with FACS sorting and flow cytometry services, and CRG Genomics Unit for assistance with sequencing services. Furthermore we would like to thank Amaya Abad and Taisia Polidori for experimental support; Beatrice Borsari, Bruna Correa and Marina Ruiz for their comments and suggestions, and Romina Garrido for administrative support.

