## Supplementary data for "Paired guide RNA CRISPR-Cas9 screening for protein-coding genes and lncRNAs involved in transdifferentiation of human B-cells to macrophages"

**Supplementary Figures: S1 to S12**

**Supplementary Tables: S1 to S11**

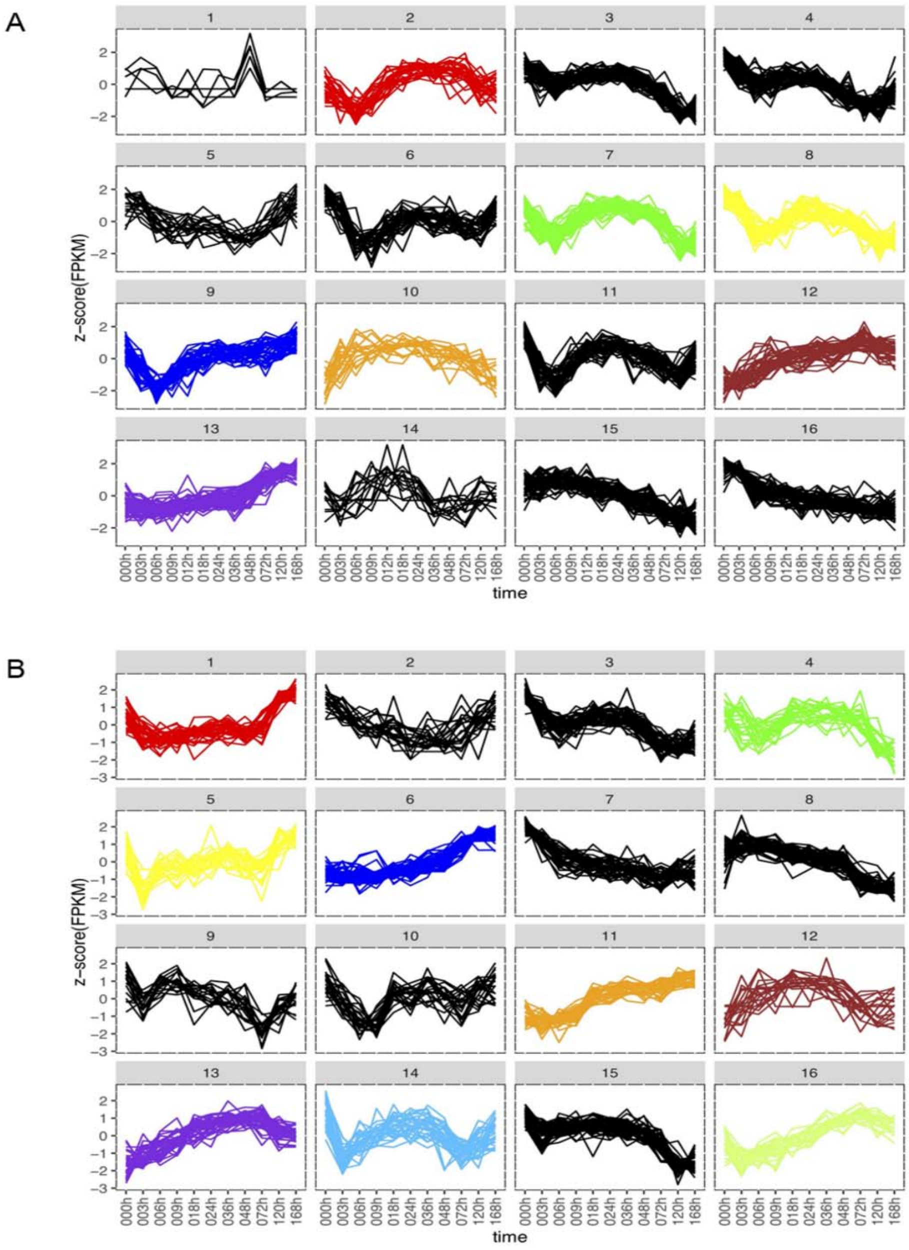


**Supplementary Figure 1: Expression clusters of lncRNAs during transdifferentiation.**

Clusters of lncRNAs with similar expression characteristics during transdifferentiation of BLaER1 cells to macrophages. lncRNAs clustered into 16 expression profiles by k-means clustering. FPKM values were log10 transformed and normalized by z-transformation. Results from 2 biological replicates are shown (**A**) and (**B**). Clusters used for the CRISPR library are highlighted using the colour that labels the genes in Figure 1C.

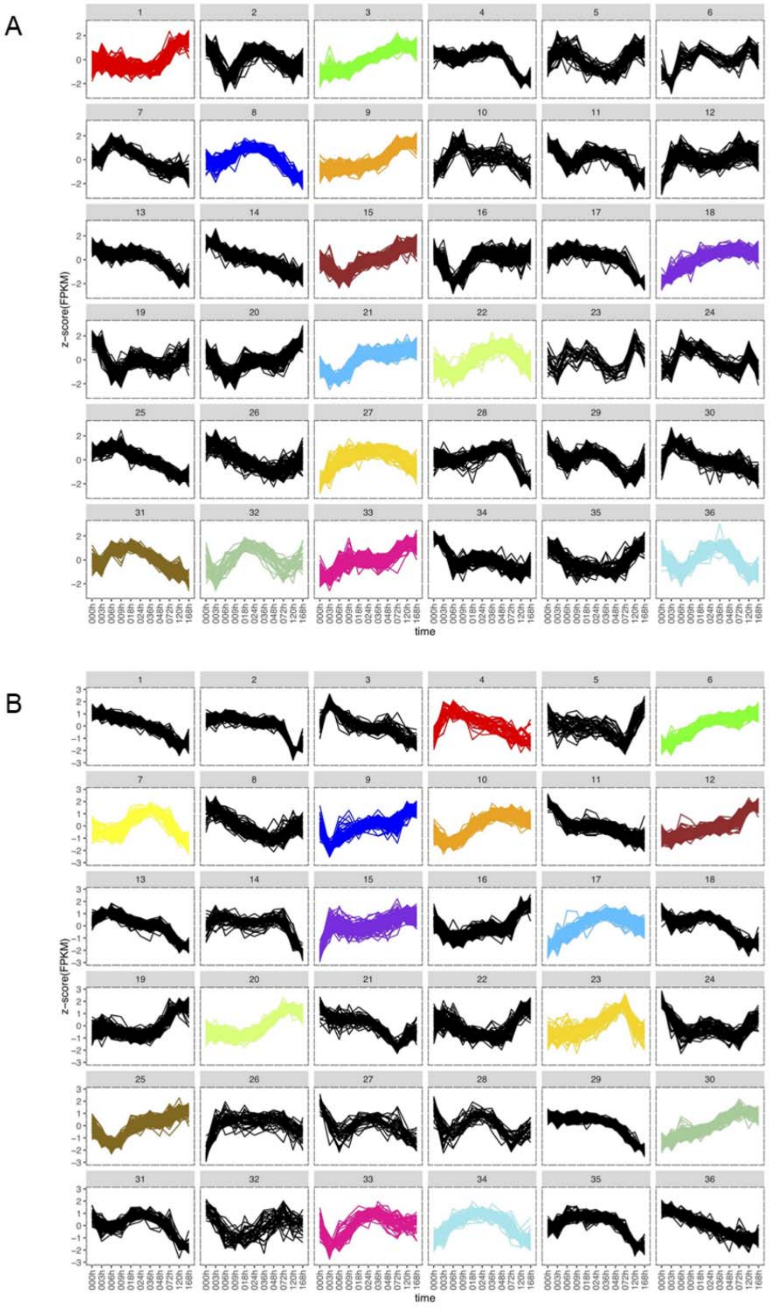


**Supplementary Figure 2: Expression clusters of protein coding genes during transdifferentiation.**

Clusters of protein coding genes with similar expression characteristics during transdifferentiation of BLaER1 cells to macrophages. Pc-genes clustered into 36 expression profiles by k-means clustering. FPKM values were log10 transformed and normalized by z-transformation. Results from 2 biological replicates are shown (**A**) and (**B**). Clusters used for the CRISPR library are highlighted using the colour that labels the genes in Figure 1C.

A

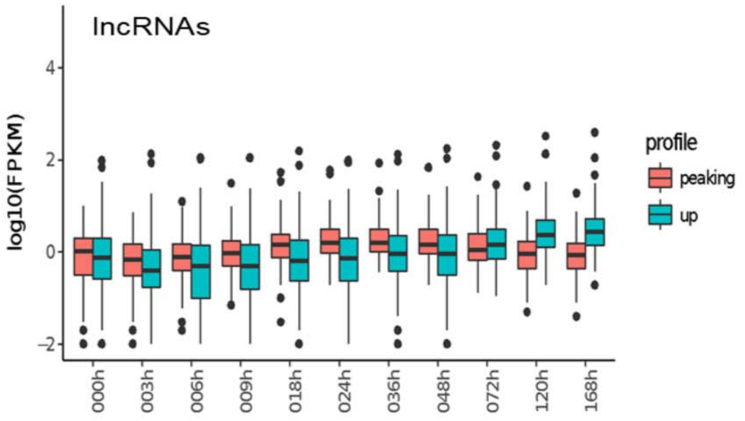


B

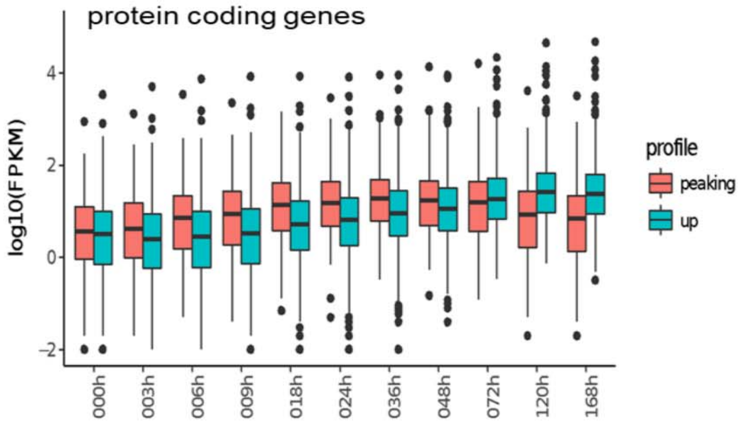


**Supplementary Figure 3**: **Expression profiles of lncRNAs and protein coding genes during transdifferentiation.**

Log10 transformed expression profiles of the 163 lncRNAs (**A**), and of the 939 protein coding genes (**B**), with peaking (red) or increasing (blue) expression.

A

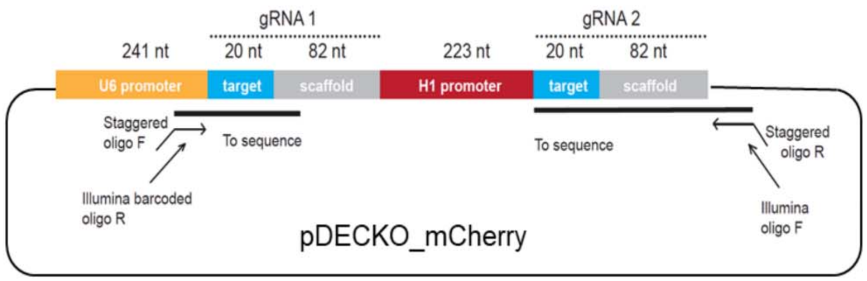


B

(Illumina oligo R )5' CAAGCAGAAGACGGCATACGAGATaaccaacgGTGACTGGAGTTCAGACGTGTGCTCTTC
(Staggered oligo F) TTCAGACGTGTGCTCTTCCGATCT(SSSSS)GTGGAAAGGACGAAACACCg
5' GTGGAAAGGACGAAACACCGGCTACAGTTAAAAAGGCCAACGTTTTAGAGCTAGAAATAGCAAGTTAA----------------3'

111 nt (sgRNA 1 site)

3' CGACTTCCAAATCATTATTTCAAAATCTCGATCTTTATCGTTCAATTTTATTCCGATCAGGCAATAGTTGAACTTTTTCACCGTGGCTCAGCCACGAAAAAAATTCGAACCGCATTGATCTAGAAC----------TCT 5'
AAATTCGAACCGCATTGATCTAGAAC(SSSSS)TCTAGCCTTCTCGCAGCACATCCC (Staggered oligo R)
CTTCTCGCAGCACATCCCTTTCTCACAagatccaaCACATCTAGAGCCACCAGCGGCATAGTAA 5' (Illumina oligo F)

200 nt (sgRNA 2 site)

Green: sgRNAs for control intergenic
Blue: scaffold sequence

**Supplementary Figure 4**: **pDECKO plasmid and sequencing oligos binding scheme.**

(**A**) Scheme of pDECKO_mCherry plasmid. H1 and U6 promoters drive the expression of the two gRNAs. The primers of the two PCR amplification steps performed before deep sequencing are also shown (see also Supplementary Tables S6 and S7). In the first PCR step, staggered oligos anneal to the U6 promoter and to the pDECKO backbone. In the second PCR step, primers containing the sample barcode and P5/P7 sequences for sequencing are added. (**B**) Oligo binding sites are shown for Illumina sequencing amplicons. The amplicon length (until the sgRNA) is 111 nt for the forward strand and 200 nt for the reverse strand. The constant scaffold sequences and the variable sgRNAs are shown in blue and green respectively. 1st PCR oligos (Staggered oligos) are shown with the staggered oligos labeled with S. 2nd PCR oligos (Illumina oligos) are shown with the barcode sequences indicated in red (see also Supplementary Tables S6 and S7).

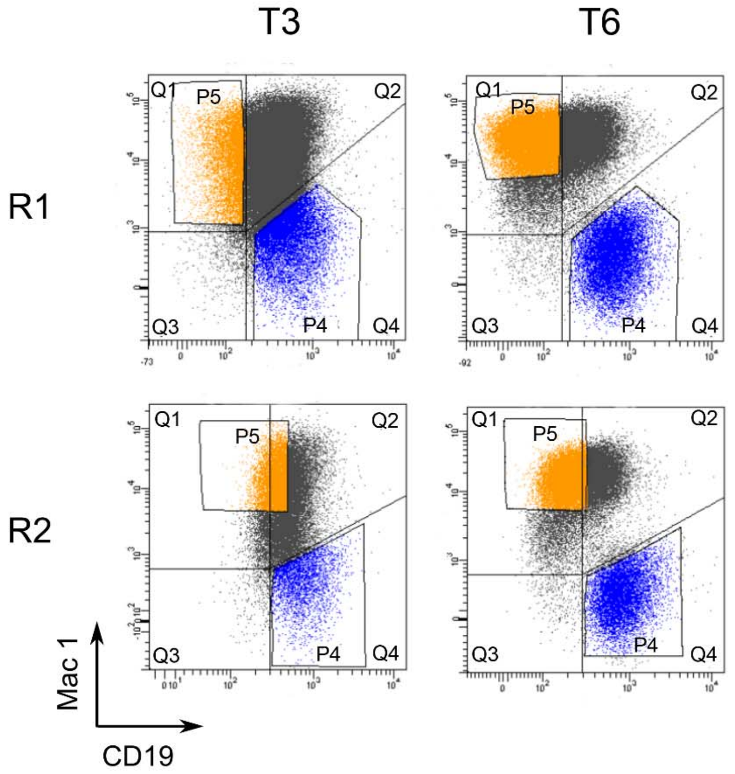


| | | Q1 | Q2 | Q3 | Q4 | P4 | P5 |
|---|---|---|---|---|---|---|---|
| R1 | T3 | 10.6% | 72.5% | 0.8% | 16% | 11.7% | 8.9% |
| | T6 | 46.1% | 39.7% | 0.5% | 13.8% | 13.4% | 38.5% |
| R2 | T3 | 12.4% | 73.8% | 1.3% | 12.5% | 10.7% | 31.3% |
| | T6 | 46.8% | 37.2% | 0.7% | 15.3% | 14.7% | 49% |

**Supplementary Figure 5: FACS sorting of BLaER1-Cas9 library.**

BLaER1-Cas9 cells infected with the pDECKO CRISPR library were transdifferentiated for T3 and T6. The cells were stained with antibodies against surface markers CD19 for B-cells and Mac-1 for macrophages, and FACS sorted. Sorted gates for cells delayed in transdifferentiation (gate P4 in blue) and cells normally differentiating (gate P5 in orange) are shown for two biological replicates (R1 and R2). The percentage of cells in each quadrant and gates is indicated.

A

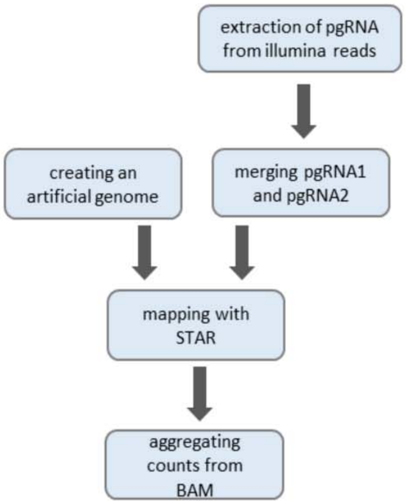


B

| | D0_total | D3_total | D6_total | D3_diff | D3_del | D6_diff | D6_del |
|---|---|---|---|---|---|---|---|
| Total reads | 23174328 | 20461721 | 17622205 | 33261946 | 35163035 | 21620659 | 28360044 |
| Uniquely mapped reads | 13648684 | 12123869 | 10320734 | 18582918 | 19804799 | 12840942 | 15755565 |
| Uniquely mapped reads % | 58.90% | 59.25% | 58.57% | 55.87% | 56.32% | 59.39% | 55.56% |
| Ambiguous reads | 120157 | 106017 | 102014 | 165894 | 195573 | 107748 | 235855 |
| Ambiguous reads % | 0.52% | 0.52% | 0.58% | 0.50% | 0.56% | 0.50% | 0.83% |
| Unmappable reads % | 40.52% | 40.15% | 40.78% | 43.55% | 43.06% | 40.03% | 43.56% |

| | D0_total | D3_total | D6_total | D3_diff | D3_del | D6_diff | D6_del |
|---|---|---|---|---|---|---|---|
| Total reads | 26644511 | 26966759 | 25395692 | 24415795 | 33691107 | 20150115 | 24517712 |
| Uniquely mapped reads | 17547956 | 17617769 | 16316787 | 15763071 | 18564557 | 12993696 | 14626243 |
| Uniquely mapped reads % | 65.86% | 65.33% | 64.25% | 64.56% | 55.10% | 64.48% | 59.66% |
| Ambiguous reads | 131929 | 138169 | 138888 | 116321 | 216878 | 99229 | 169568 |
| Ambiguous reads % | 0.50% | 0.51% | 0.55% | 0.48% | 0.64% | 0.49% | 0.69% |
| Unmappable reads % | 33.55% | 34.06% | 35.10% | 34.82% | 44.15% | 34.90% | 39.55% |

C

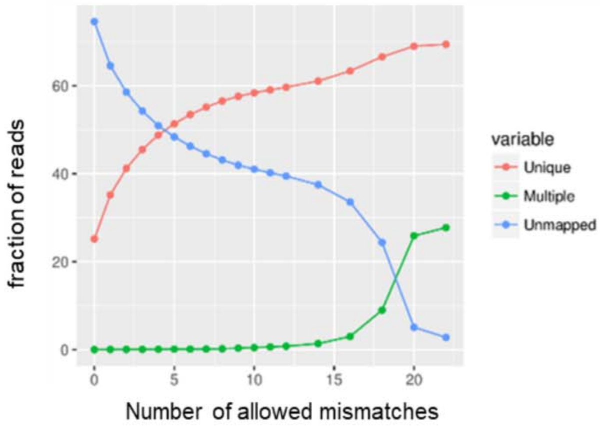


D

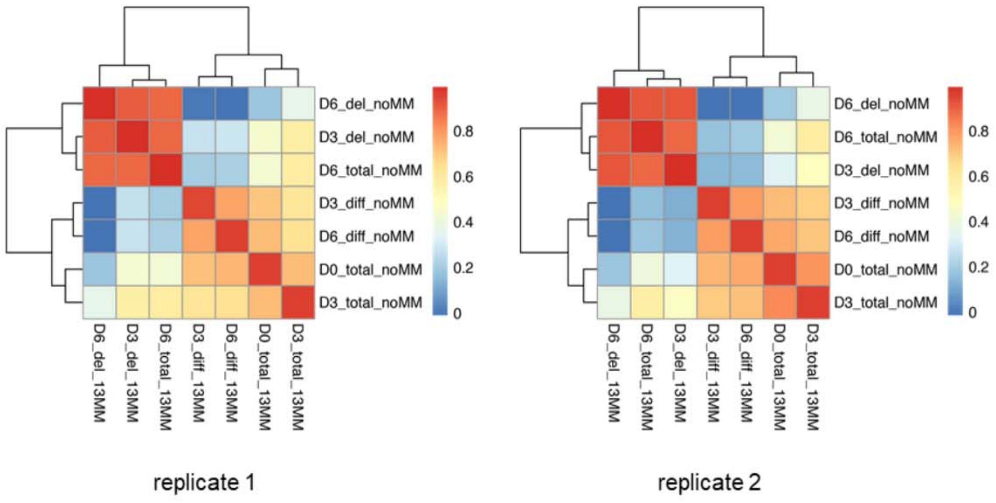

**Supplementary Figure 6: Statistics on quantification of pgRNA representation in the screening.**

(**A**) Schematic flow diagram displaying the steps from the pgRNA FASTQ sequences after sequencing to count tables per pgRNA. In short, pgRNA sequences are extracted from FASTQ sequences by finding the proximal constant plasmid sequence. pgRNA2 is reverse complemented (only needed for paired end sequencing) and merged with pgRNA1. Both are mapped as one sequence to the merged expected sequences converted into artificial chromosomes with STAR mapper. Count tables are generated from BAM files by aggregation. (**B**) Detailed mapping statistics for both replicates in the quantification. Initial read counts per sample ranged between 20 to 35 million reads of which on average 55% can be mapped against perfect library sequences with not more than 13 mismatches (diff = differentiated population, del = population with delayed differentiation). (**C**) Uniquely mapped, multi-mapped and unmapped reads as a function of allowed mismatches during quantification of the sequenced samples in a range of 0 to 26 mismatches. (**D**) Pearson correlations of guide pair quantification between runs allowing for up to 13 mismatches and only allowing perfect matches. Correlation values for identical source samples were ranging between 0.95 and 1. (diff = differentiated cell population from FACS orange gate P5, del = cell population with delayed differentiation from FACS blue gate P4). Results for two biological replicates are shown.

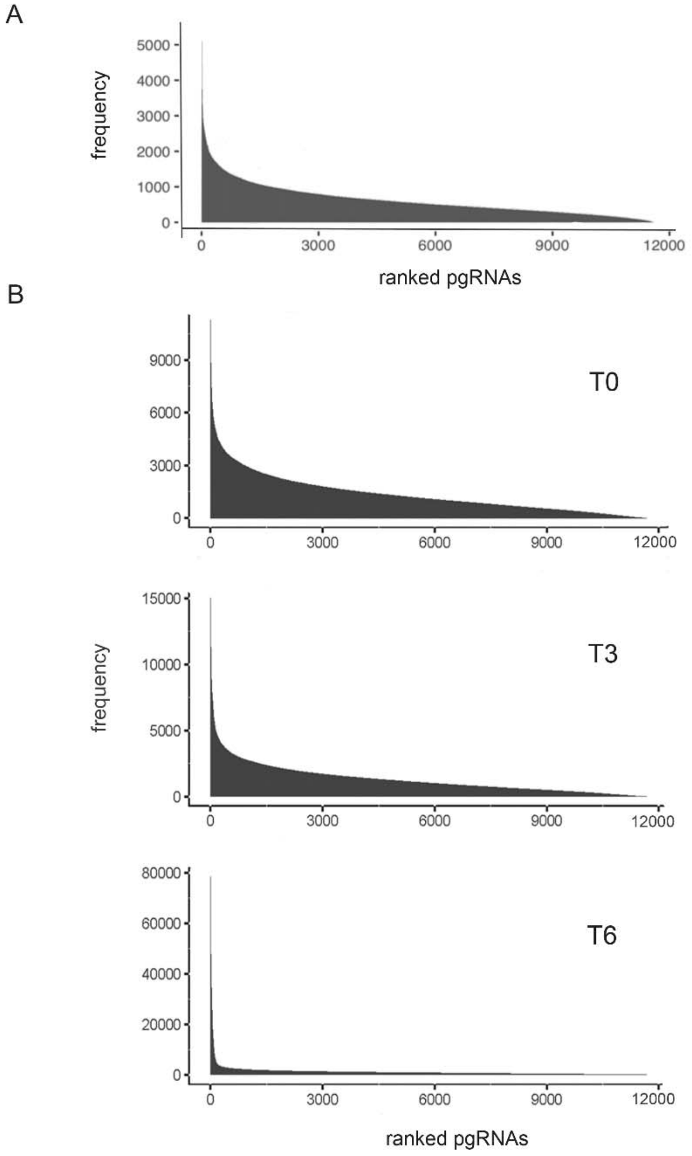


**Supplementary Figure 7: Quantification of pgRNA distribution before and during screening.**

(**A**) Ranked distribution of counts per pgRNA in the initial library (after cloning). The library showed a good representation and homogeneity (> 95 % of coverage and a delta of top to bottom decile < 10 fold, respectively), as described in (3). (**B**) Ranked distribution of counts per pgRNA in the control samples at T0, T3 and T6 upon transdifferentiation induction, that contain all cells independent of B-cell and macrophage marker abundance.

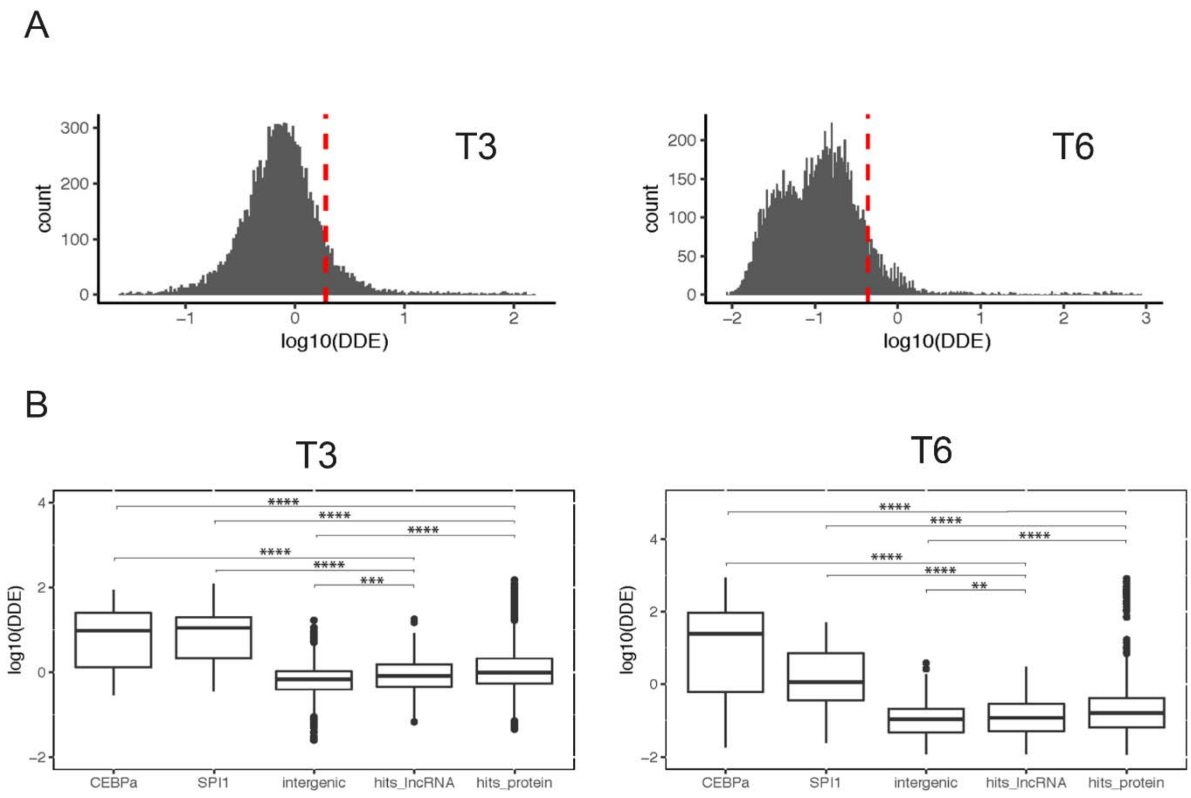


**Supplementary Figure 8: Target genes disrupted by CRISPR-Cas9.**

(**A**) Distribution of DDE values at T3 and T6 upon transdifferentiation induction. DDE is computed as the ratio of normalized counts from the delayed subpopulation (del) divided by the counts from the transdifferentiated population (dif). Highest decile is marked by a dashed red line. Values are the average between the two biological replicates. (**B**) Comparison of DDE values for the selected candidate pgRNAs targeting lncRNAs and protein coding genes of the highest decile (from panel A) at T3 and T6 upon transdifferentiation induction, with CEBPa and SPI1 as positive controls, and intergenic regions as negative controls. Values are the average between the two biological replicates. Significant values were assessed by means of the Wilcoxon test.

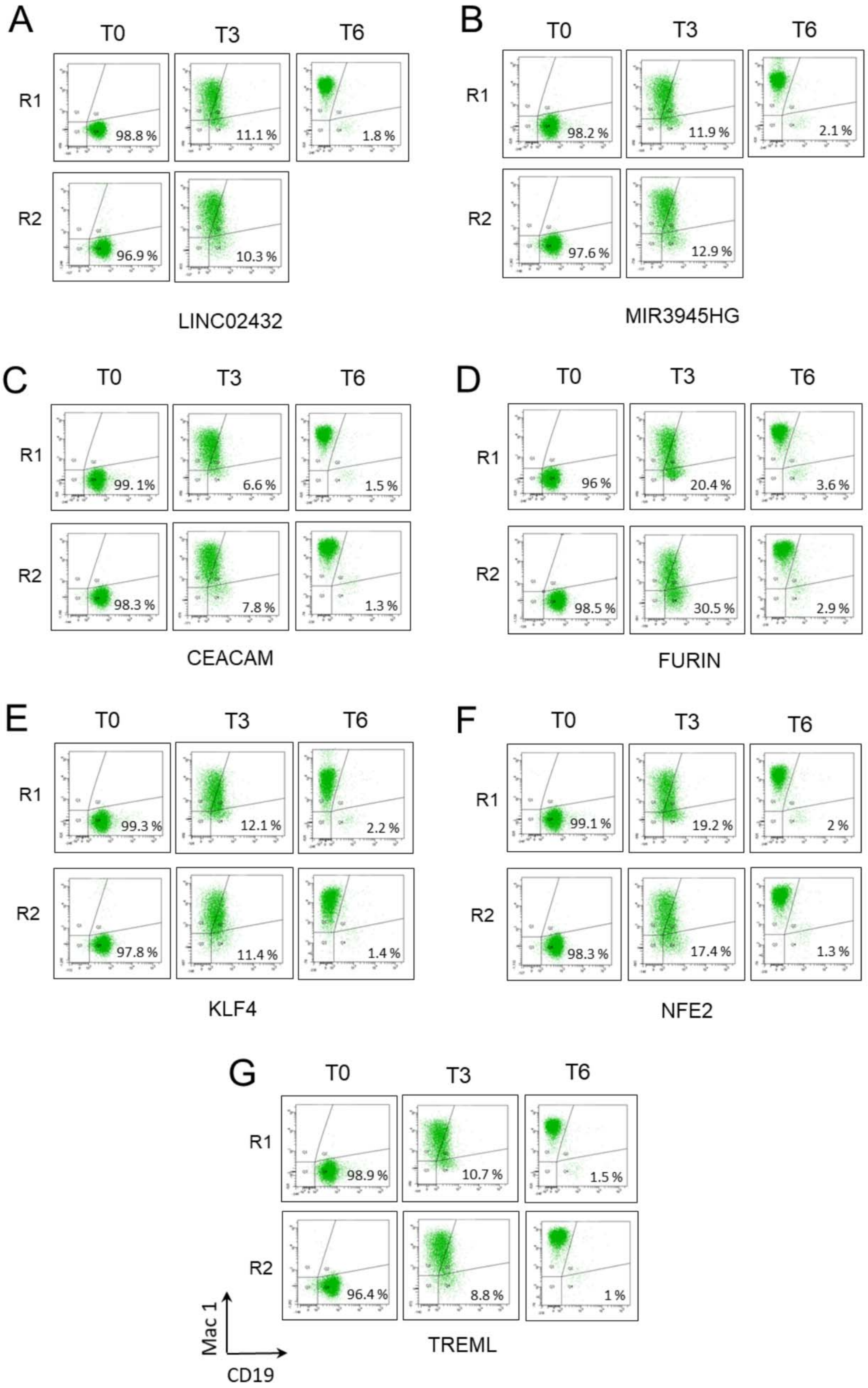
A
T0
T3
T6
R1
98.8 %
11.1 %
1.8 %
R2
96.9 %
10.3 %
LINC02432
B
T0
T3
T6
R1
98.2 %
11.9 %
2.1 %
R2
97.6 %
12.9 %
MIR3945HG
C
T0
T3
T6
R1
99. 1%
6.6 %
1.5 %
R2
98.3 %
7.8 %
1.3 %
CEACAM
D
T0
T3
T6
R1
96 %
20.4 %
3.6 %
R2
98.5 %
30.5 %
2.9 %
FURIN
E
T0
T3
T6
R1
99.3 %
12.1 %
2.2 %
R2
97.8 %
11.4 %
1.4 %
KLF4
F
T0
T3
T6
R1
99.1 %
19.2 %
2 %
R2
98.3 %
17.4 %
1.3 %
NFE2
G
T0
T3
T6
R1
98.9 %
10.7 %
1.5 %
R2
96.4 %
8.8 %
1 %
Mac 1
CD19
TREML

**Supplementary Figure 9: Individual target validation by flow cytometry.**

Flow cytometry analysis of targets validated from the screening at 0, 3, and 6 days (T0, T3 and T6) of transdifferentiation. CD19 was used as a B-cell marker on the X-axis and Mac-1 was used as a macrophage marker on the Y-axis. Cells that do not undergo transdifferentiation (delayed fraction) remain in quadrant Q4 (percentages of cells for this quadrant are shown at T3 and T6 upon transdifferentiation induction). (**A**) LINC02432; (**B**) MIR3945HG; (**C**) CEACAM ; (**D**) FURIN; (**E**) KLF4; (**F**) NFE2; (**G**) TREML2. Two biological replicates (R1 and R2) are shown except for LINC02432 and MIR3945HG from which only one biological replicate was available at T6.

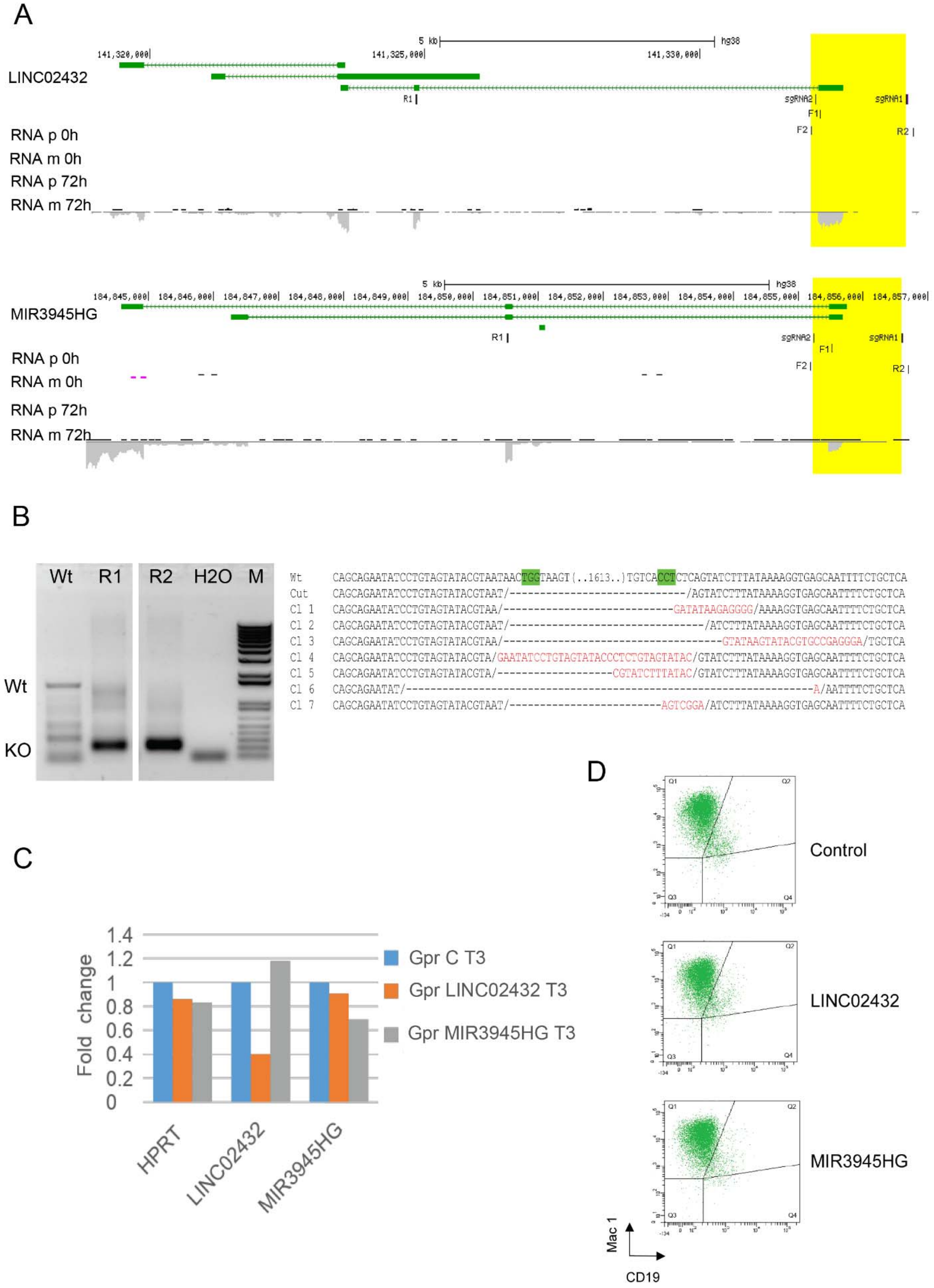
A
LINC02432
5 kb
hg38
141,320,000
141,325,000
141,330,000
R1
sgRNA2
sgRNA1
F1
F2
R2
RNA p 0h
RNA m 0h
RNA p 72h
RNA m 72h
MIR3945HG
184,845,000
184,846,000
184,847,000
184,848,000
184,849,000
184,850,000
184,851,000
184,852,000
184,853,000
184,854,000
184,855,000
184,856,000
184,857,000
B
Wt
R1
R2
H2O
M
Wt
KO
Cut
Cl 1
Cl 2
Cl 3
Cl 4
Cl 5
Cl 6
Cl 7
C
Fold change
1.4
1.2
1
0.8
0.6
0.4
0.2
0
HPRT
LINC02432
MIR3945HG
Gpr C T3
Gpr LINC02432 T3
Gpr MIR3945HG T3
D
Control
LINC02432
MIR3945HG
Mac 1
CD19

**Supplementary Figure 10: lncRNA target sites and individual validations.**

(**A**) Diagrams from UCSC genome browser showing the binding sites of pgRNAs (in yellow) for *LINC02432* (ENSG00000248810.1) and *MIR3945HG* (ENSG00000251230.1). qPCR primers binding sites are indicated as F1/R1 and primers for checking CRISPR cut are indicated as F2/R2. RNA seq expression is shown at 0h and 72h of transdifferentiation. (**B**) Left, agarose gel showing the PCR products from genomic DNA amplification of wild type cells (wt) and the knockout band (KO) of *LINC02432* CRISPR-Cas9 edited cells (from FACS sorted enriched cell population) for two biological replicates (R1 and R2), a water template (H20) was used as a PCR negative control. Wild type and knockout bands have an expected size of 1817 nt and 179 nt respectively. Right, Sanger sequencing results from clones obtained with TA cloning method (PAM sequences are shown in green, insertions are shown in red and deletions are shown as dashed lines). (**C**) qRT-PCR to check the expression of the lncRNAs *LINC02432* and *MIR3945HG* after GapmeR treatment and cell differentiation for 3 days (T3). The plot shows expression of the housekeeping gene HPRT, and of the two lncRNAs in cells treated with each of the three GapmeRs. (**D**) Flow cytometry analysis at T3 of differentiation of BLaER cells treated with GapmeR control (against an intergenic region), or GapmeRs against *LINC02432* and *MIR3945HG*.

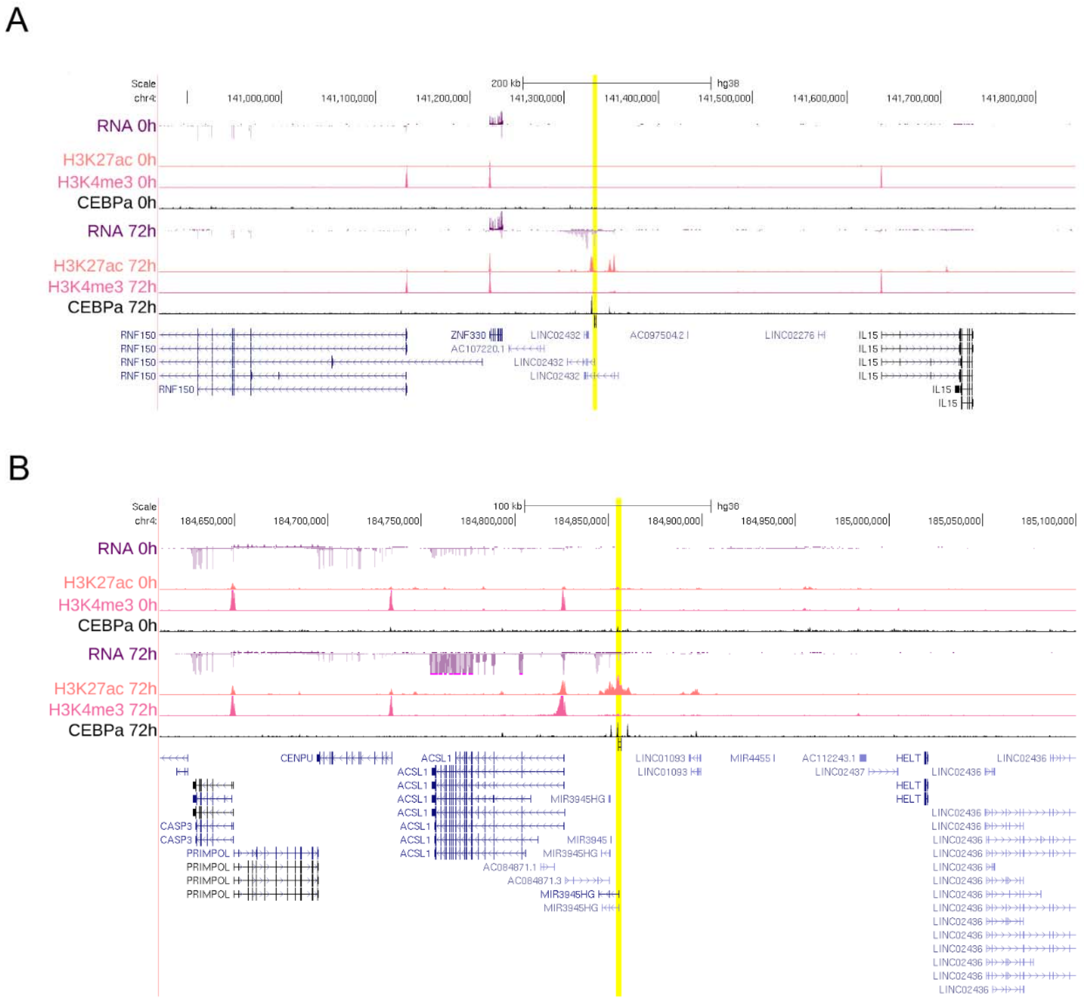


**Supplementary Figure 11: Epigenetic landscape of the candidate lncRNAs.**

UCSC genome browser diagrams for LINC02432 (**A**) and MIR3945HG (**B**). pgRNAs binding sites are highlighted in yellow. In both cases, there is an increase of H3K27ac mark, associated with active promoters and enhancers, matching the binding of CEBPa and the upregulation of the lncRNAs expression at 72h after induction. However, there is no increase in H3K4me3 mark, typically associated with active promoters. This is consistent with these regions behaving as enhancers.

A

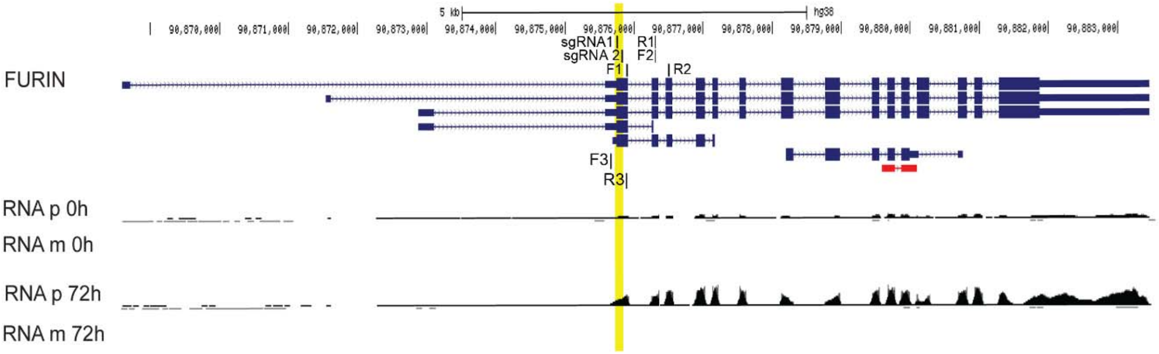


B

Wt CCCTGGTTGCTATGGGTGGTAGCAGCAACAGGAACCTTGGTCCTGCTAGCAGCTGATGCTCAGGGCCAGAAGGTCTTCACCAACACGTGGGCTGTGCGCAT

KO CCCTGGTTGCTAT/ ------------------------------------------------------------/ACGTGGGCTGTGCGCAT

cl 1 CCCTGGTTGCTAT/ ------------------------------ TTGCTATAGGACCCC ------------------------------/ACGTGGGCTGTGCGCAT

cl 2 CCCTGGTTGCTAT/ ------------------------------ TTGCTAT- GG ------------------------------/ACGTGGGCTGTGCGCAT

cl 3 CCCTGGTTGCTAATGGGTGGTAGCAGCAACAGGAACCTTAGTCCTGGTCCTGCTAGCAGCTGATGCTCAGGGCCAGAAGGTCTTCACCAACCGGACGTGGGCTGTGCGCAT

cl 4 CCCTGGTTGCTAGGGCTGGGTGGTAGCAGCAACAGGAACCTTGGTCCTGCTAGCAGCTGATGCTCAGGGCCGGAAGGTCTTCACCAACCCACGTGGGCTGTGCGCAT

cl 5 CCCTGGTTGCTAGAGACGGCGCCGTCGTGCTCCCCGGAGTGGAGGCGGCCGAGCGCGCCGGGGTGCCCGCCTTCCTGGAGACCTCCGCGCCCCGCAACCTCCCCTTCTACGAGGCTGTGCGCAT

cl 6 CTGNTTGCC/------------------------------AACCTTGGTCCTGCTAGCAGCTGATGCTCAGGGCCAGAAGGTCTTCACCAAC----GTGGGCTGTGCGCAT

cl 7 CCCTGGTTGCTATGCCGGGGTGGTAGCAGCAACAGGAACCTTGGTCCTGCTAGCAGCTGATGCTCAGGGCCAGAAGGTCTTCACCAAGGGCCGTGGGCTGTGCGCAT

C

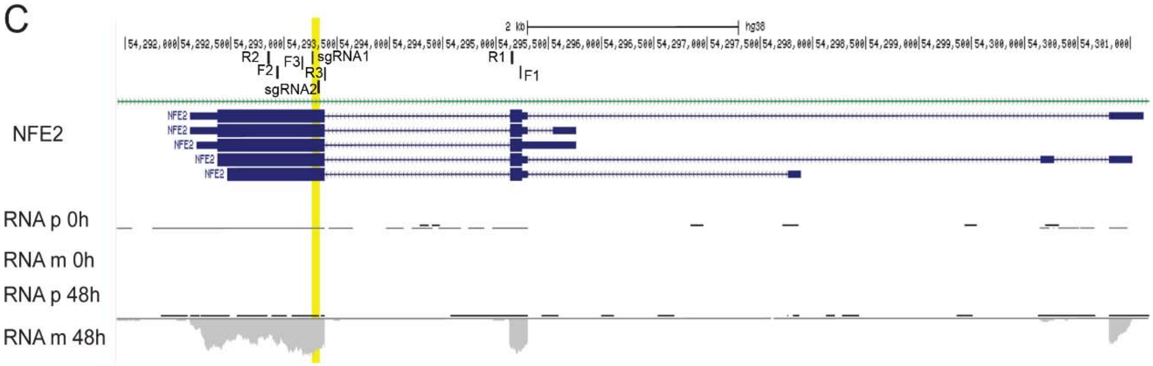


D

Wt CAGCTCCATACCTTGGACCTCCACCACCCACAACTTACTGCCCCTGCTCAATCCACCCAGATTCTGGCTTCCCACTTCCT

KO CAGCTCCATACCTTGGACC/------------------------------/TTCTGGCTTCCCACTTCCT

cl1 NGCTGCCATACCTTGGACCTCCACCACCCACAACTTACTGCCCCTGCTCAATCCACCCAG/-----------/GCTTCCCACTTCCT

cl2 CAGC----CA-----GCTTGGA-NTCCACCGCCCACAACTTACTGCCCCTGCTCAATCCACCCAGATTCTGGCTTCCCACTTCCT

cl3 CAGCT------TACCTTGGACCTCCACCACCCACAACTTACTGCCCCTGCTCAATCCACCCAGAGTCTTTCTGGCTTCCCACTTCCT

cl4 TACCTTGGACCTCCACCACCCACAACTTACTGCCCCTGCTCAATCCACCCAGAGAGTGTTCGTTTCTGGCTTCCCACTTCCT

**Supplementary Figure 12: FURIN and NFE2 target sites and TA cloning.**

UCSC genome browser diagrams for *FURIN* (**A**) and *NFE2* (**C**). pgRNAs binding sites are highlighted in yellow. The two primer pairs used for qRT-PCR are indicated as F1/R1 and F2/R2. Primers used for checking CRISPR deletion are indicated as F3/R3. Sanger sequencing results are shown from clones obtained with TA cloning: (**B**) for pDECKO-FURIN infected cells and (**D**) for pDECKO-NFE2 infected cells. PAM sequences are shown in green, insertions are shown in red and deletions are shown as dashed lines.

### SUPPLEMENTARY TABLES

**Supplementary Table S1:** gRNA sequences.

**Supplementary Table S2:** CRISPETa output lists for controls and targets including final CRISPR 165 oligonucleotide library.

**Supplementary Table S3**: Oligos for pgRNA cloning in pDECKO_Cherry.

**Supplementary Table S4:** Oligos for sgRNA cloning of CD19 in pDECKO_mCherry.

**Supplementary Table S5:** List of cloned plasmids.

**Supplementary Table S6:** Sequences of the staggered oligos.

**Supplementary Table S7:** Sequences of Illumina oligos.

**Supplementary Table S8:** Sequences of oligos for qRT-PCR.

**Supplementary Table S9:** Sequences of oligos outside the CRISPR cut region.

**Supplementary Table S10:** Shortlisted candidates from screening.

**Supplementary Table S11:** RNA seq expression data of protein coding genes and lncRNA final targets during BLaER1 transdifferentiation.
